# Molecular Determinants of Functional Bacterial sRNA–mRNA Interactions Revealed by Integrating RNA Interactomes and Interpretable Machine Learning

**DOI:** 10.64898/2026.08.26.747384

**Authors:** Fatemeh Safari, Daniel G. Mediati, Saleh Alquethamy, Jai J Tree, Fatemeh Vafaee

**Affiliations:** School of Biotechnology and Biomolecular Sciences, University of New South Wales (UNSW Sydney), Kensington, NSW, 2052, Australia; UNSW AI Institute, UNSW Sydney, Kensington, NSW, 2052, Australia; Australian Institute for Microbiology and Infection, University of Technology Sydney, Sydney, New South Wales, Australia; ARC Centre of Excellence in Synthetic Biology, Department of Natural Sciences, Macquarie University, Sydney, New South Wales, Australia

## Abstract

Bacterial small RNAs (sRNAs) regulate gene expression by base pairing with target mRNAs, yet transcriptome-wide interactome mapping has shown that many sRNA–mRNA interactions detected *in vivo* have modest or no regulatory effect using orthogonal reporter assays. The features that determine functional outcome remain poorly defined. Here, we integrated Hfq-CLASH interactome mapping with matched transcriptomic and proteomic profiling in *Escherichia coli* and developed an interpretable machine-learning framework to identify the determinants that distinguish functional from non-functional interactions. Using sequence, structural, thermodynamic, duplex and protein-occupancy features, transcriptomic and proteomic responses were predicted with above-chance performance, achieving AUCs of 0.78 and 0.74, respectively. Feature attribution revealed that physical pairing alone is insufficient for regulation; instead, regulatory outcome is shaped by a coordinated interplay between RNA secondary structure, thermodynamic accessibility and local protein-binding context. Target-side Hfq occupancy emerged as a positive predictor of functional regulation, whereas AR2-domain occupancy on the sRNA was associated with non-responsive interactions, suggesting that distinct ribonucleoprotein states may separate productive regulation from non-productive binding. These findings indicate that the regulatory fate of an sRNA–mRNA interaction is an emergent property of its biophysical context and protein-binding environment, rather than a direct consequence of physical pairing alone.

**GRAPHICAL ABSTRACT:** 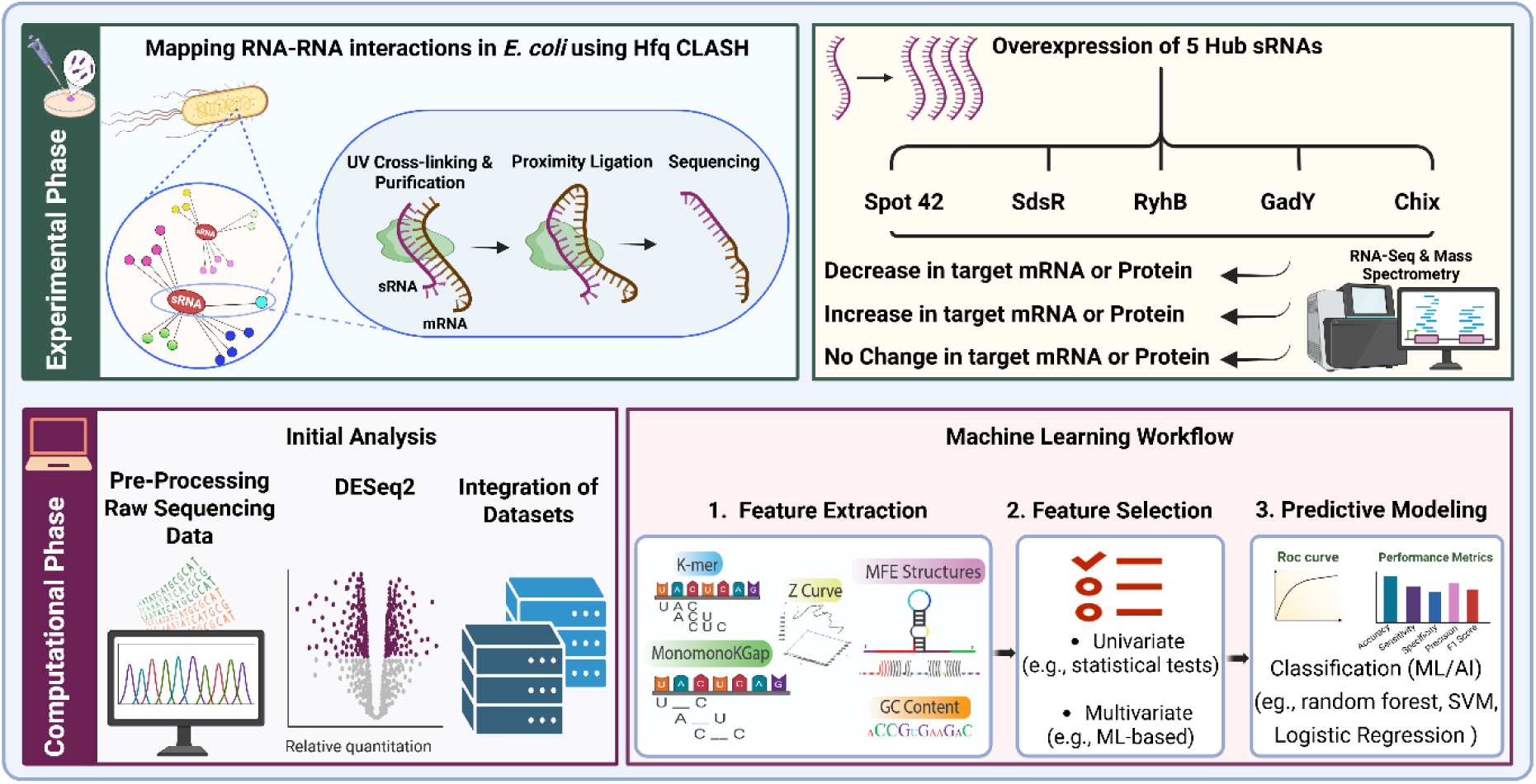

## INTRODUCTION

Bacterial survival depends on the ability to rapidly and precisely adapt gene expression in response to fluctuating environmental conditions, nutrient availability, host interactions, and cellular stress (1). While transcriptional regulation has long been considered the primary mechanism of gene control, it is now clear that post-transcriptional regulation constitutes an equally important layer of gene expression management (2,3). Central to this layer are small regulatory RNAs (sRNAs), a diverse class of non-coding RNAs, typically 50–500 nucleotides in length, that modulate gene expression primarily through complementary base pairing with target transcripts (4–6). In pathogenic bacteria, this regulatory capacity is closely linked to host adaptation and virulence, with sRNAs contributing to the control of invasion, intracellular survival, stress tolerance, immune interaction, and virulence-associated gene expression, and thereby influencing infection outcome (7–10). Defining the molecular rules that determine sRNA function is therefore central both to bacterial physiology and to antimicrobial discovery and RNA-based biotechnology.

RNA base pairing is governed not only by complementarity, but also by the accessibility, position and structural context of the interacting regions (8,11). Many bacterial sRNAs contain modular RNA elements, including a structured 3′ terminator that contributes to stability, an RNA chaperone-binding region, and a conserved base-pairing domain that often functions through short seed-like interactions with target mRNAs. In some cases, target recognition is initiated through exposed nucleotides in stem-loop regions, followed by additional base-pair formation and structural rearrangement, whereas other sRNAs pair through single-stranded regions that enable rapid recognition of complementary mRNA sites (8,12). In many bacteria, RNA chaperones such as Hfq facilitate this process by binding both sRNAs and mRNAs, promoting RNA annealing, stabilising regulatory RNAs, and influencing whether base pairing results in translational repression, activation or altered mRNA degradation (13). Thus, functional sRNA targeting reflects a coordinated interplay between sequence motifs, structural accessibility, and recruitment of RNA-binding proteins and ribonucleases.

Mechanistic understanding of bacterial sRNA regulation has largely emerged from low-throughput studies of individual sRNA–mRNA pairs, which cannot explain the principles governing transcriptome-wide regulatory networks. Approaches such as CLASH (cross- linking, ligation and sequencing of hybrids), RIL-seq (RNA interaction by ligation and sequencing) and related proximity-ligation methods now detect RNA–RNA interactions *in vivo* at transcriptome scale, revealing extensive bacterial RNA interactomes and many previously unrecognised sRNA–mRNA interactions (14,15). However, these advances have also exposed a major conceptual gap: while physical interactions can now be identified at unprecedented scale, the determinants that distinguish functional from non-functional interactions remain largely unresolved. This challenge became particularly clear in studies that combined RIL-seq interactome mapping with functional profiling through sRNA overexpression (16). These analyses revealed a striking dichotomy, in which only a discrete subset of physically bound mRNA targets showed significant expression changes, whereas many verified interactions remained functionally unaffected. This finding established that physical pairing, although necessary, is not sufficient to predict regulatory outcome. Instead, target response was closely associated with differences in Hfq occupancy, suggesting that competition for the RNA chaperone is a critical determinant of regulatory function, beyond base-pairing potential alone (16). Therefore, a central unresolved challenge in bacterial RNA biology is to define, in a quantitative and mechanistically interpretable manner, the molecular features that determine whether an sRNA–mRNA interaction produces a functional regulatory outcome.

Parallel advances in transcriptomics and proteomics provide an opportunity to move from interaction discovery toward functional interpretation. Transcriptomic profiling following sRNA perturbation captures regulatory consequences at the RNA level, including changes in transcript abundance, stability and RNA decay, while proteomic profiling can reveal downstream effects on protein abundance that may arise through altered translation or post-transcriptional regulation (17). Examining these molecular layers is therefore essential for identifying the molecular features that contribute to functional outcomes, rather than simply cataloguing physical sRNA-mRNA interactions. This distinction is particularly important because transcriptome-wide interaction mapping studies have shown that many experimentally detected sRNA-mRNA interactions do not necessarily produce measurable changes in target expression, indicating that physical pairing alone is insufficient to define regulatory function (14,18).

Addressing this challenge requires a framework that can identify the local molecular context that determines regulatory outcome. In this study, local context refers to the sequence, structural, thermodynamic and protein-binding properties surrounding each sRNA-mRNA interaction. These include RNA composition, base-pairing features, predicted structural accessibility, duplex energetics and experimentally derived occupancy signals from RNA-binding proteins such as Hfq and RNase E. Such features are particularly relevant because protein-RNA interaction sites can contextualise sRNA-mRNA pairing and may help explain why some interactions produce transcriptomic or proteomic changes whereas others remain functionally silent.

Distinguishing functional from non-functional sRNA–mRNA interactions therefore depends on integrating these diverse, high-dimensional molecular features. Machine learning is well suited to this task because it can identify predictive patterns across multiple interacting feature classes (19,20). In RNA biology, feature extraction is a critical step that transforms raw RNA sequence and structural information into quantitative representations suitable for modelling, including nucleotide composition, k-mer patterns, structural accessibility, folding properties and interaction energetics (21). In the context of bacterial sRNA regulation, such features can be combined with experimentally derived protein occupancy signals, including Hfq and RNase-associated binding information, to investigate how local RNA-RNA and protein-RNA context contributes to transcriptomic and proteomic outcomes.

Here, we present an interpretable machine-learning framework that integrates experimentally mapped RNA interactomes, transcriptomic and proteomic responses, and protein-occupancy profiles to identify the molecular features that distinguish functional sRNA–mRNA interactions from functionally silent pairs detected *in vivo*. Rather than focusing solely on interaction prediction, our approach provides mechanistic insight into how sequence, RNA structure, thermodynamic accessibility and protein-binding context collectively shape bacterial post-transcriptional regulation.

## MATERIAL AND METHODS

### Experimental Procedures

*Bacterial Strains and Growth Conditions. E. coli* str. MG1655 Δ*chiX*, Δ*gadY*, Δ*ryhB*, Δ*sdsR*, and Δ*spot42* strains were generated by λ Red recombination and verified by PCR and Sanger sequencing. Each strain carried either the corresponding L-arabinose-inducible pBAD+1 complementation plasmid or an empty-vector control. Strains were grown in LB medium at 37 °C with shaking at 220 rpm and supplemented with ampicillin (100 µg mL⁻¹). *sRNA Overexpression and Transcriptome Profiling. E. coli* MG1655 strains carrying individual deletions of *chiX*, *gadY*, *ryhB*, *sdsR*, or *spot42* were transformed with either an empty pBAD+1 or the corresponding sRNA complementation plasmid. Strains were cultured in LB medium supplemented with ampicillin 100 µg.mL^-1^ at 37°C with shaking at 220 RPM. Three independent biological replicates were prepped for each strain. Cultures were grown to mid-exponential phase (OD600 = 0.6), and sRNA expression was induced with 0.2% (w/v) L-arabinose for 30 min. Following induction, cultures were immediately treated with 95:5 (v/v) ethanol:phenol stop solution. Cells were harvested and mechanically lysed with zirconia beads using a FastPrep-24 5G bead-beating system. Total RNA was then isolated using GTC-phenol extraction. Residual genomic DNA was removed by treatment with RQ1-RNase-free DNase (Promega). RNA concentration was determined using a Qubit fluorometer, and RNA integrity was assessed using a TapeStation (Agilent).

RNA samples were submitted to Novogene for directional prokaryotic RNA-sequencing library preparation, including depletion of bacterial rRNA. Libraries were sequenced using a 150-bp paired-end strategy on an Illumina NovaSeq platform.

### Hfq-CLASH for In Vivo Interactome Mapping

CLASH was performed as previously described (22) with some modifications specific for *E. coli*. MG1655 WT and Hfq-HTF (dual-affinity-tagged HTF strain) (23). Detailed methods are described in the Supplementary Methods.

### Protein extraction and Mass Spectrometry (LC-MS/MS)

Cultures for proteomic analysis were grown in parallel with the RNA-seq cultures on the same day and under identical growth and induction conditions. Cultures were harvested by centrifugation, and cell pellets were resuspended in lysis buffer containing 50 mM Tris–HCl (pH 7.8; Sigma-Aldrich), 150 mM NaCl, 0.1% (v/v) NP-40, and cOmplete EDTA-free protease inhibitor (one tablet per 50 mL; Roche). Cells were disrupted with zirconia beads using a FastPrep-24 5G homogenizer (MP Biomedicals) for two 40-s cycles separated by a 1-min rest. Lysates were clarified by centrifugation at 5,000 × g for 10 min, and the resulting supernatants were centrifuged at 16,000 × g for 20 min. Protein concentrations were determined using the Bio-Rad DC Protein Assay and normalised across samples. Samples were submitted to the UNSW Bioanalytical Mass Spectrometry Facility for label-free quantitative LC–MS/MS analysis using an Orbitrap Fusion Lumos Tribrid mass spectrometer (Thermo Fisher Scientific).

Raw LC–MS/MS data were processed by the UNSW Bioanalytical Mass Spectrometry Facility using MaxQuant (24). Tandem mass spectra were searched against the UniProt *E.coli* K-12 MG1655 reference proteome (25), and protein abundance was determined using label-free quantification (LFQ) method. The mass spectrometry proteomics data have been deposited to the ProteomeXchange Consortium via the PRIDE (26) partner repository with the dataset identifier PXD082677.

### Computational Analysis and Predictive Modelling

#### Overview of the Computational Framework

We developed a computational framework to predict the functional outcomes of sRNA–mRNA interactions and to identify the molecular features associated with regulatory activity. The framework integrates three complementary data sources: (i) Hfq-CLASH data providing direct evidence of in vivo RNA–RNA interactions, (ii) CRAC (UV cross-linking and analysis of cDNAs) data quantifying protein occupancy on RNA transcripts (27), and (iii) functional readouts derived from RNA-seq and mass spectrometry-based proteomics datasets. For each sRNA–mRNA interaction pair, a comprehensive set of quantitative molecular features was extracted to characterise sequence composition, physicochemical properties, RNA secondary structure, interaction thermodynamics, predicted duplex architecture, and experimental protein-binding context. These features were used as inputs for supervised machine learning pipelines designed to model functional outcomes independently across three regulatory layers: transcriptomic response, proteomic response, and interaction abundance.

### Hfq-CLASH Data Processing and Interaction Set Curation

RNA-RNA interactions were recovered from Hfq-CLASH sequencing dataset using the Hyb-CRAC-R pipeline (28), generating a comprehensive interactome comprising 9,742 unique RNA–RNA interactions. To focus the analysis on biologically well-characterised regulatory interactions, this global set was filtered to retain interactions in which one partner corresponded to one of the five studied sRNAs (ChiX, GadY, RyhB, SdsR or Spot42) and the other partner corresponded to a protein-coding mRNA. The resulting curated sRNA-mRNA interaction set formed the core dataset for downstream transcriptomics-level, proteomics-level and interaction-level modelling. Hfq-CLASH sequencing data is deposited at NCBI GEO under accession GSE343154.

### Transcript Annotation and Coordinate Resolution

A subset of interaction entries in the initial CLASH dataset contained incomplete or ambiguous annotations, including entries labelled as “unknown”. To resolve these cases, a coordinate- and strand-aware re-annotation pipeline was developed using 5′ and 3′ untranslated region (UTR) coordinate data from the UTR_5_3_sequence.tsv file from RegulonDB (29) as a reference. This reference dataset provides UTR start and stop positions, transcription unit information, strand orientation, and flanking gene assignments. UTR coordinates were parsed into numeric start and stop positions, and strand labels were standardised as “+” and “−”.

For each hybrid arm, entries annotated as unknown were tested for positional overlap with annotated UTR intervals using a ±10 nucleotide tolerance window. If an overlap occurred within a 5′ UTR, the *first_gene* was assigned for entries on the “+” strand, whereas the *last_gene* was assigned for entries on the “−” strand. Conversely, if an overlap occurred within a 3′ UTR, the *last_gene* was assigned for entries on the “+” strand, whereas the *first_gene* was assigned for entries on the “−” strand. The RNA class was updated to “5′ UTR” or “3′ UTR” accordingly. Entries without positional overlap with annotated UTR intervals remained classified as unknown. This process reduced the fraction of unlabelled hybrids and improved the gene-level interpretability of the interaction dataset.

### Protein Occupancy Quantification from CRAC Data

To characterise protein-binding context within interaction regions, strand-specific occupancy metrics were derived from CRAC read-depth data for three proteins: full-length RNase E, the AR2 domain of RNase E, and the RNA chaperone Hfq. UV-crosslinking of RNase E and AR2 are described in (30). For each protein, bedGraph files from three biological replicates were processed separately for plus- and minus-strand data. For every sRNA–mRNA interaction pair, genomic coordinates and strand information were extracted for both the sRNA and mRNA partners. Within each interaction region, the maximum read depth across overlapping intervals was identified in each replicate, and replicate-level maxima were averaged to generate a robust per-interaction occupancy estimate. Regions with no detectable CRAC signal were assigned an occupancy value of zero. This analysis generated six protein-occupancy features per interaction pair: maximum read depth on the sRNA and mRNA regions for RNase E, AR2, and Hfq. These features provide quantitative measures of local protein occupancy that may influence RNA accessibility, duplex formation, directed RNA decay, or regulatory outcome. RNase E-CRAC and CRAC data for the AR2 sub-domain have been deposited at NCBI GEO under the accession GSE317719.

### RNA-seq Analysis and Transcriptomic Functional Classification

Functional outcomes at the transcriptomic level were determined using differential expression analysis of RNA-seq data. Differential expression analysis was performed using READemption (31). RNA sequencing datasets for sRNA deletion mutants and pulse expression strains are available at NCBI GEO under the accession number GSE342672.

### Feature Extraction and Molecular Characterisation

To enable machine learning analysis, each curated sRNA–mRNA interaction was represented as a quantitative feature vector describing its molecular properties. As illustrated in Figure 1, the feature extraction workflow integrated sequence composition, RNA secondary structure, thermodynamic properties, protein occupancy and intermolecular duplex characteristics to capture both the individual RNA partners and their local interaction context. In total, 824 features were extracted for each interaction and organised into the categories summarised in Table 1. Detailed descriptions of the feature extraction strategies and their computational rationale are provided in our comprehensive review of RNA feature extraction methods (21)

**Figure 1.**
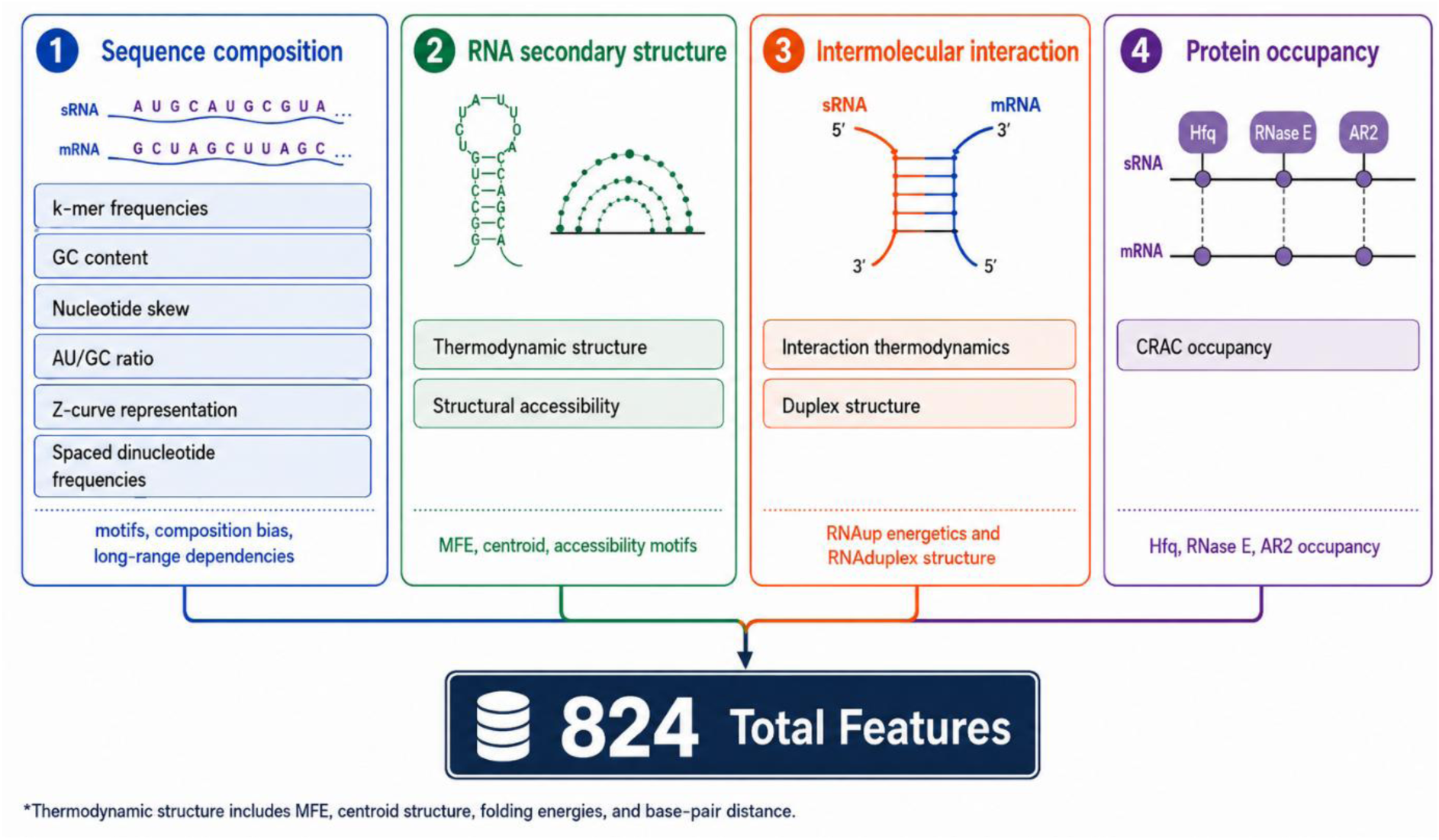
Molecular feature encoding framework for sRNA–mRNA interaction prediction. Schematic overview of the 824 descriptors extracted from each curated sRNA–mRNA pair for machine learning analysis. Features were grouped into four major categories: sequence composition, RNA secondary structure, intermolecular interaction, and protein occupancy. Sequence composition features included k-mer frequencies, GC content, nucleotide skew, AU/GC ratio, Z-curve descriptors, and spaced dinucleotide frequencies. Structural features captured thermodynamic properties and structural accessibility, whereas interaction features described binding energetics and duplex architecture. Protein occupancy features represented maximum binding signals of Hfq, RNase E, and AR2 across the interaction regions. The infographic was generated using GPT Image 2 and subsequently reviewed by the authors for scientific accuracy.

**Table 1.** Molecular feature categories extracted for each sRNA–mRNA interaction. The 824 descriptors comprise sequence-composition, RNA secondary-structure, intermolecular-interaction and protein-occupancy features.

| **Feature Category** | **Feature Type** | **Description** | **No. of Features** |
| --- | --- | --- | --- |
| **Sequence composition** | k-mer frequencies | Frequencies of all possible 3-mers (64) and 4-mers (256) calculated separately for sRNA and mRNA sequences to capture local sequence motifs. | 640 |
|  | GC content | Percentage of G and C nucleotides for each RNA. | 2 |
|  | Nucleotide skew | GC skew and AU skew describing nucleotide compositional asymmetry. | 4 |
|  | AU/GC ratio | Ratio of AU to GC nucleotides for each RNA. | 2 |
|  | Z-curve representation | Three normalized descriptors (x, y, z) representing nucleotide composition in 3D space for each RNA. | 6 |
|  | Spaced dinucleotide frequencies | Frequencies of dinucleotide pairs separated by gaps of 1, 2 or 3 nucleotides (monoMonoKGap) to capture long-range sequence dependencies. | 96 |
| **RNA secondary structure** | Thermodynamic structure | Minimum free energy (MFE), centroid structure energy and centroid distance calculated using ViennaRNA. | 6 |
|  | Structural accessibility | Frequencies of structural 3-mer and 4-mer motifs derived from paired/unpaired (P/U) secondary structure representation. | 48 |
| **Intermolecular interaction** | Interaction thermodynamics | RNAup-derived duplex energetics including hybridization energy, unfolding penalties for sRNA and mRNA, and total interaction energy. | 4 |
|  | Duplex structure | RNA duplex-derived descriptors including interaction length, base-pair composition, mismatches, bulges and longest consecutive base-paired region. | 9 |
|  | Total hybrid count | Total number of hybrid reads supporting each sRNA–mRNA interaction, providing a measure of the experimental representation of the interaction in the CLASH dataset. | 1 |
| ***Protein occupancy*** | CRAC occupancy | Maximum occupancy of Hfq, RNase E and AR2 over the sRNA and mRNA interaction regions. | 6 |
| ***Total*** |  | **All sequence, structural, thermodynamic, duplex and protein-occupancy descriptors used for machine learning.** | **824** |

### Predictive Modelling Framework

Three independent supervised classification pipelines were developed to predict sRNA– mRNA interaction outcomes across three complementary biological readouts: transcriptomic response, proteomic response and total hybrid-count abundance. Although all pipelines used the same broad feature space, including sequence-derived, structural, thermodynamic and protein-occupancy descriptors, each was tailored to the sample size, class distribution and label definition of its corresponding dataset.

The pipelines are described separately below because they address distinct classification tasks. However, they were built around three shared design principles. First, all preprocessing, feature selection and model fitting were performed strictly within the training partitions. Second, model interpretation was carried out using SHAP values and standardised logistic-regression coefficients, allowing each retained feature to be assigned both a relative contribution and a direction of association with the outcome. Third, the resampling and class-balancing strategy was selected according to the statistical structure of each dataset.

For the transcriptomics pipeline, the dataset was strongly imbalanced, with 59 responsive and 380 non-responsive interactions. Stratified cross-validation alone would therefore place only a small number of minority-class samples in each test fold, resulting in unstable performance estimates. To address this, we used majority-class subsampling combined with bootstrap resampling, which retained all minority-class samples while evaluating the model across multiple non-overlapping representations of the majority class. In contrast, the proteomics dataset was near-balanced, with 127 responsive and 111 non-responsive interactions, while the hybrid-count dataset showed only moderate imbalance, with 126 high-count and 313 low-count interactions. For these two pipelines, repeated stratified three-fold cross-validation preserved class proportions across folds and provided robust variance estimates without discarding data; therefore, additional subsampling was not required.

### Transcriptomics Pipeline: Predicting mRNA-Level Regulatory Response

The transcriptomics pipeline classified each sRNA–mRNA interaction by whether the target mRNA responded to sRNA overexpression, using a pre-specified |log₂FC| ≥ 0.5 threshold used in bacterial sRNA-focused transcriptomic and interactome studies (32,33). Of the 439 curated Hfq-CLASH interactions, 59 (13.4%) were classified as responsive (class 1) and 380 (86.6%) as non-responsive (class 0), a class imbalance of approximately 6.4:1. We defined responsiveness by effect size rather than by adjusted-significance filters, which are underpowered for the modest, fine-tuned shifts characteristic of sRNA regulation and would systematically reclassify genuinely regulated targets into the majority class. The full modelling workflow is shown in Figure 2.

**Figure 2.**
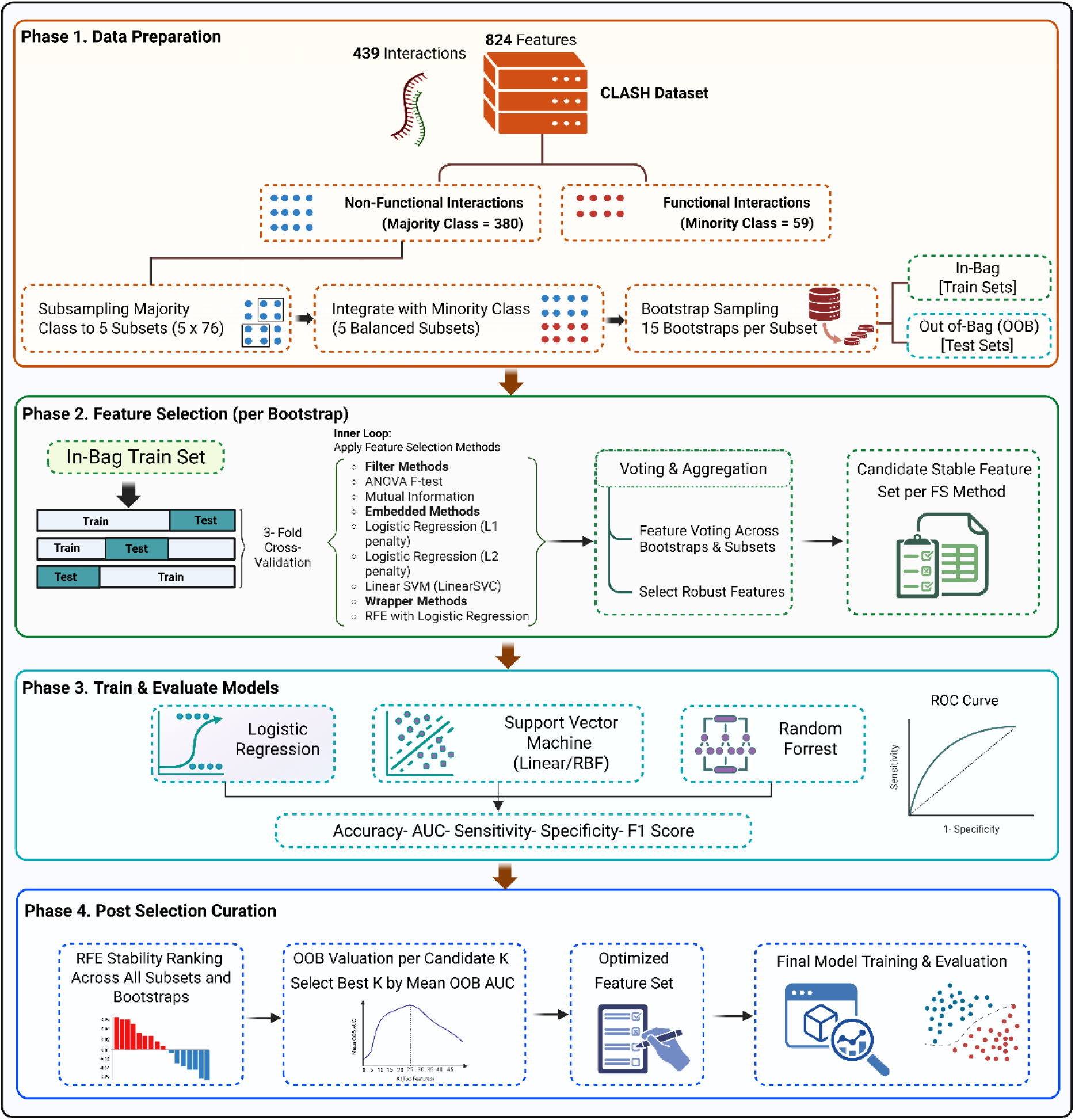
Machine learning pipeline for classifying sRNA-mRNA interactions as functional or non-functional based on target mRNA expression changes after sRNA overexpression. Created with BioRender.

### Class-Imbalance Handling: Majority-Class Subsampling with Bootstrapping

To address the strong class imbalance in the transcriptomics dataset without using synthetic oversampling, a majority-class subsampling strategy was applied. The 380 non-functional interactions were divided into five non-overlapping subsets of 76 samples, ensuring that each majority-class sample appeared in one subset. Each subset was then combined with all 59 functional interactions, generating five partially balanced datasets of 135 samples each. Within each dataset, bootstrap resampling with replacement was used to generate training sets, and samples not selected during bootstrapping were retained as out-of-bag test samples. This procedure was repeated 15 times per subset, resulting in 75 evaluation rounds in total. This design ensures that each model is trained on a dataset with improved class balance while preserving all available information from the minority class. At the same time, it allows the model to be evaluated across multiple representations of the majority class, providing a more robust estimate of generalisation performance.

### Two-Stage Feature Selection

Feature selection was performed independently within the in-bag training partition of each bootstrap resample, using a two-stage procedure repeated across all 75 evaluation rounds.

### Stage 1: Domain-aware univariate filtering

The first stage used domain-aware univariate filtering to reduce the feature space before multivariate selection. Filtering was performed separately within predefined feature groups to avoid dominance by large feature blocks, particularly k-mer features. General molecular features were assessed using ANOVA F-tests with false-discovery-rate control. For k-mer features, a stricter criterion was applied by combining FDR correction with a minimum effect-size threshold and correlation-based redundancy pruning, retaining only the top-ranked k-mers in each round. Structural k-mer and spaced dinucleotide features were filtered using the same block-wise logic. Features that did not pass this stage were excluded from subsequent multivariate feature selection.

### Stage 2: Model-based feature selection and stability voting

Following Stage 1 filtering, model-based feature selection methods were applied in parallel: L2-regularised logistic regression, L1-regularised logistic regression, linear SVM, recursive feature elimination with logistic regression, and mutual information. Each method was implemented within an inner three-fold cross-validation framework, and selected features were recorded for every bootstrap iteration across all five majority-class subsets.

Feature stability was then assessed using a two-level voting procedure. First, within each majority-class subset, a feature was retained if it was selected in at least 7 of the 15 bootstrap iterations. Second, features were consolidated across the five subsets, with features retained if they appeared in at least one subset. This generated a frozen feature list for each selection method.

### Recursive Feature Elimination for Dimensionality Reduction

To further reduce dimensionality while preserving predictive performance, recursive feature elimination (RFE) was applied to the consensus-selected features using logistic regression as the base estimator. RFE was performed within the same five-subset, 15-bootstrap out-of-bag (OOB) framework to ensure leakage-free evaluation. At each iteration, features were ranked according to the absolute magnitude of their logistic regression coefficients, and the least important feature was removed. Feature stability was then aggregated across all 75 bootstrap rounds. Candidate feature-set sizes of 10, 20, 25, 30, 35, 40, 45, 50, 55, 60, 80 and 100 were evaluated using the mean OOB area under the receiver operating characteristic curve (AUC-ROC). The feature-set size with the highest mean OOB AUC-ROC was selected as the final optimised feature panel.

### Final Model Evaluation

In the final stage, each frozen feature set was evaluated using the same subset partitions and bootstrap seeds used during feature selection, ensuring direct alignment between selection and model assessment. For each bootstrap round, models were trained on the in-bag samples using only the selected features and evaluated on the corresponding out-of-bag samples, with no additional feature selection performed. Feature standardisation was fitted exclusively on the in-bag training data and then applied to the out-of-bag test data. Multiple classifiers were evaluated, including L1- and L2-regularised logistic regression, linear and RBF SVMs, and random forest. Performance was assessed primarily using AUC, with accuracy, recall, specificity and F1 score also reported. Out-of-bag predictions were additionally aggregated at the interaction level by averaging predicted probabilities across bootstrap evaluations, providing a more stable estimate of model performance.

### Post-Hoc Model Interpretation

Model interpretation was conducted using three complementary approaches applied to the final feature list. First, true SHAP (Shapley Additive exPlanations) values were computed using the exact linear explainer across all 75 bootstrap rounds, producing interaction-level feature attribution maps. Second, standardised logistic regression coefficients were extracted from each fitted model and averaged, providing directionality of association with functional response.

### Proteomics Pipeline: Predicting Protein-Level Regulatory Response

Protein-level responsiveness was defined using a pre-specified |log₂FC|≥0.5 threshold, a biologically meaningful cutoff used in bacterial sRNA-associated studies (34,35). This yielded a balanced class distribution of 127 (53.4%) responsive and 111 (46.6%) non-responsive interactions out of 238 matched pairs. We utilized an effect-size threshold rather than statistical significance to directly capture the presence of a regulatory response. Relying on adjusted-significance filtering can be overly conservative for moderate post-transcriptional changes, which risks systematically misclassifying genuinely but modestly regulated targets as non-responsive and biasing the dataset against the subtle regulatory events the model is designed to detect.

### Cross-Validation Design

A 20-repeat stratified three-fold cross-validation scheme was employed, generating 60 total train–test splits (Figure 3). In each fold, approximately 159 samples were allocated to training and 79 to testing, with class proportions preserved by stratification. The use of 20 repeats with different random partitions provides robust variance estimates and reduces the sensitivity of performance metrics to any single train–test partition.

**Figure 3.**
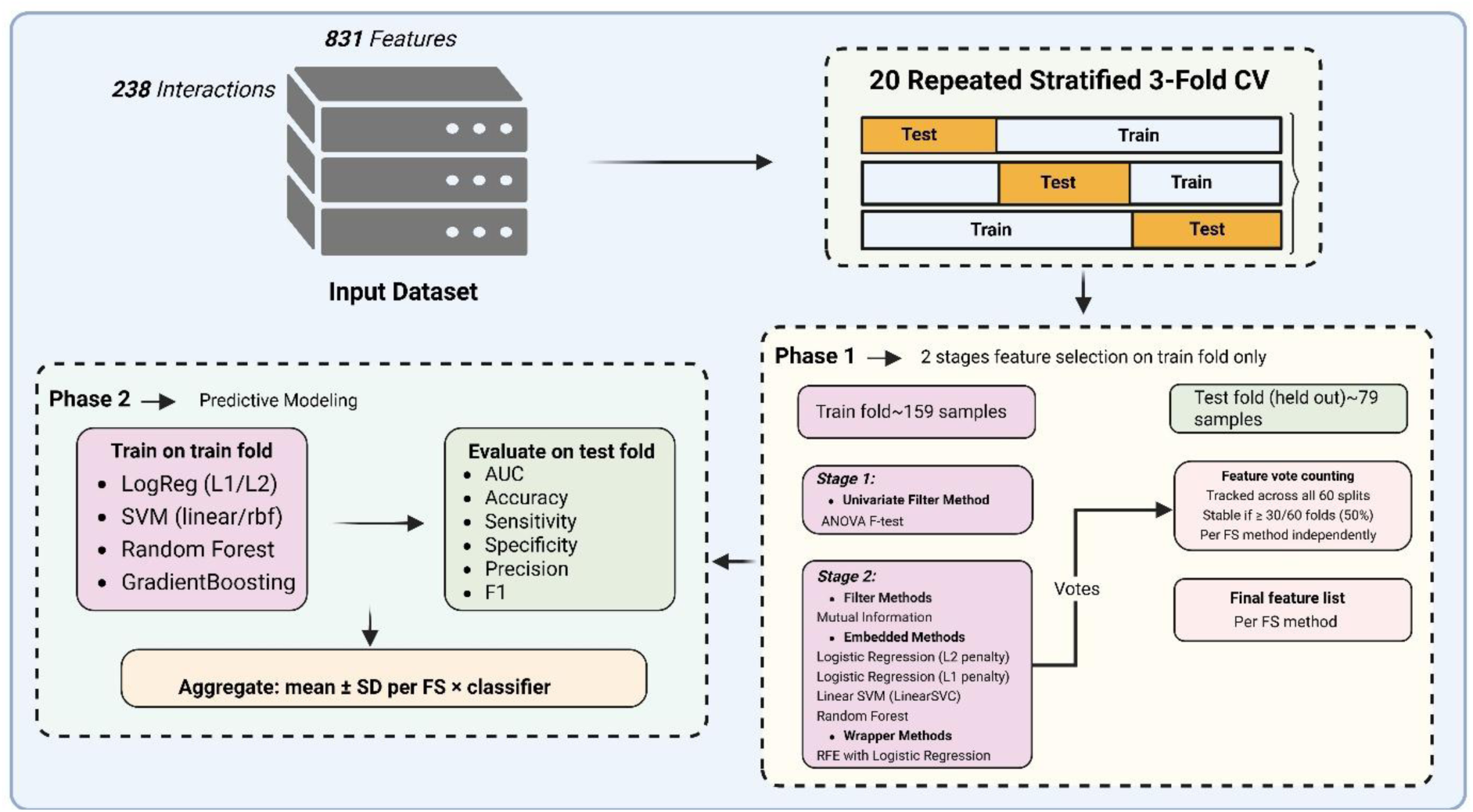
Machine learning pipeline for classifying sRNA-mRNA interactions as functional or non-functional based on target protein expression changes after sRNA overexpression. Created with BioRender.

### Two-Stage Feature Selection

Feature selection for the proteomics pipeline followed the same two-stage framework described for the transcriptomics analysis. Briefly, all preprocessing and feature-selection steps were performed independently within each of the 60 training folds, with the corresponding test folds kept completely unseen. The initial feature space was first reduced using block-wise univariate filtering across k-mer, spaced dinucleotide, structural k-mer and remaining molecular feature groups, thereby limiting the dominance of large feature classes and excluding weak candidates before model-based selection.

The filtered feature set was then subjected to multiple embedded and wrapper selection methods, including L1- and L2-regularised logistic regression, linear SVM, recursive feature elimination with logistic regression, mutual information and random forest importance. Features were standardised using parameters estimated from the training fold only. Feature stability was assessed across the 60 folds; features selected in at least 30 folds were considered method-stable, and those retained by at least two selection methods were included in the final consensus feature set.

### Final Model Evaluation and Post-Hoc Interpretation

Five classifiers were evaluated on the final feature sets using the same 20×3-fold cross-validation framework: LR-L2, LR-L1, SVM with RBF kernel, random forest, and gradient boosting.

Post-hoc interpretation followed the same framework as the transcriptomics pipeline. True SHAP values were computed using the exact linear explainer across all 60 folds, and standardised logistic regression coefficients were extracted for directionality analysis.

### Hybrid Count Pipeline: Predicting Interaction Abundance

The hybrid count pipeline classified interactions by chimeric read abundance in Hfq-CLASH, defining high-abundance interactions as total hybrid count > 5 (class 1; n = 126) and low-abundance interactions as total hybrid count ≤ 5 (class 0; n = 313). The same feature-encoding framework was used as in the other pipelines, except that total hybrid count was removed from the predictor set because it defined the outcome label, leaving 823 features for modelling.

The total hybrid count pipeline followed the same modelling structure as the proteomics pipeline (Figure 4). Briefly, a 20-repeat stratified three-fold cross-validation design was used, generating 60 train-test splits. Within each training fold, the same two-stage feature-selection framework was applied, comprising ANOVA-based univariate filtering followed by multi-method feature selection using LR-L1, LR-L2, random forest, mutual information, RFE-LR and LinearSVM. Feature stability was assessed across the 60 folds, and all preprocessing steps were fitted on the training data only before being applied to the held-out test fold. Model performance was evaluated across the same classifier panel and summarised using AUC, accuracy, sensitivity, specificity, and F1 score. Model interpretation followed the same post hoc framework, using SHAP values and standardised logistic regression coefficients.

**Figure 4.**
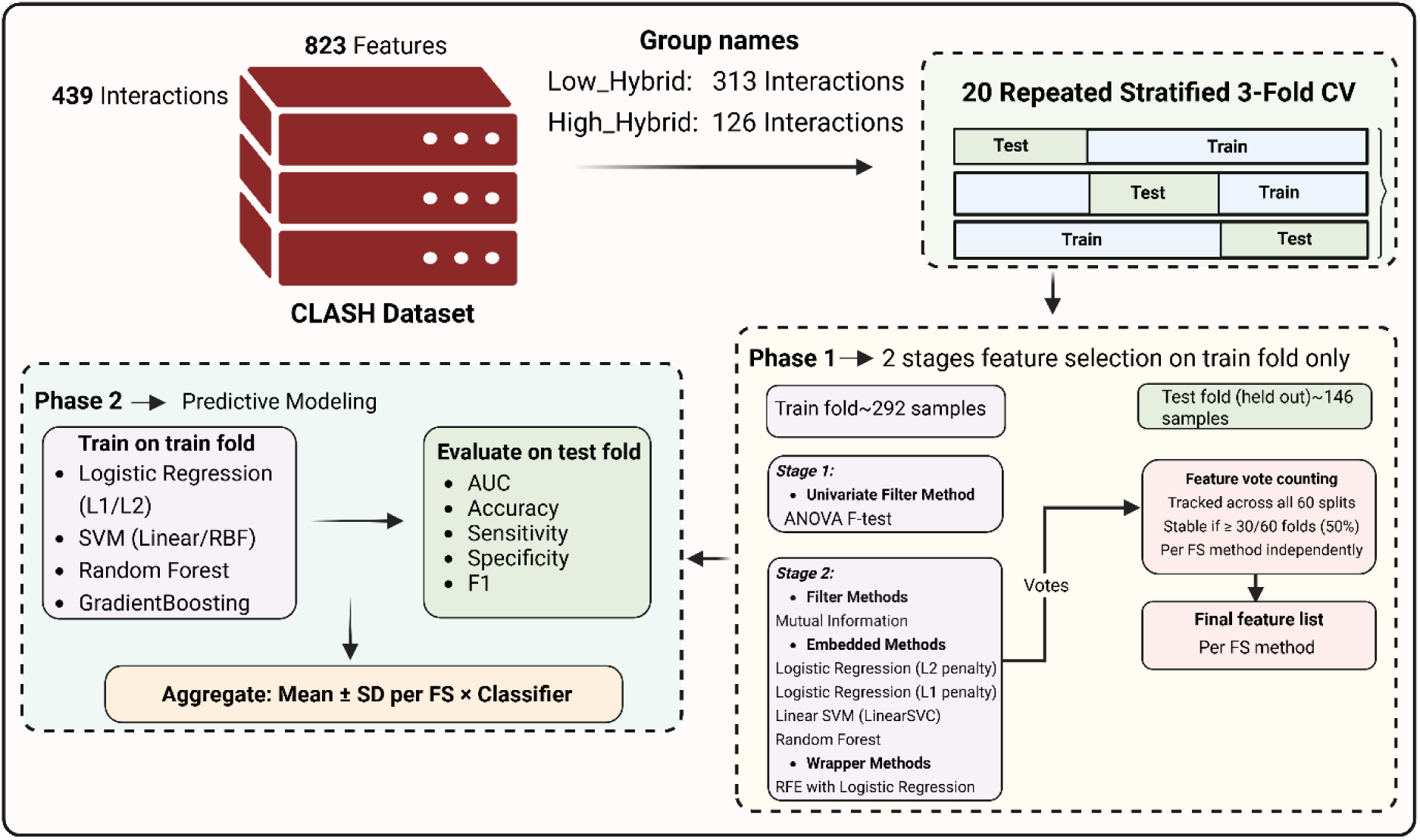
Overview of the hybrid-count modelling workflow for predicting sRNA–mRNA interaction abundance. Created with BioRender.

## RESULTS

### Mapping the sRNA-mRNA interactome and defining functional outcomes

To link physical sRNA–mRNA interactions with regulatory consequences in *E. coli*, we integrated Hfq-CLASH interaction mapping with matched transcriptomic and proteomic profiling following pulsed overexpression of five hub sRNAs in their isogenic deletion background: ChiX, GadY, RyhB, SdsR and Spot42. From 9,742 global RNA–RNA interactions, we curated 439 sRNA–mRNA pairs involving these five sRNAs and protein-coding targets (Table 2).

**Table 2.**
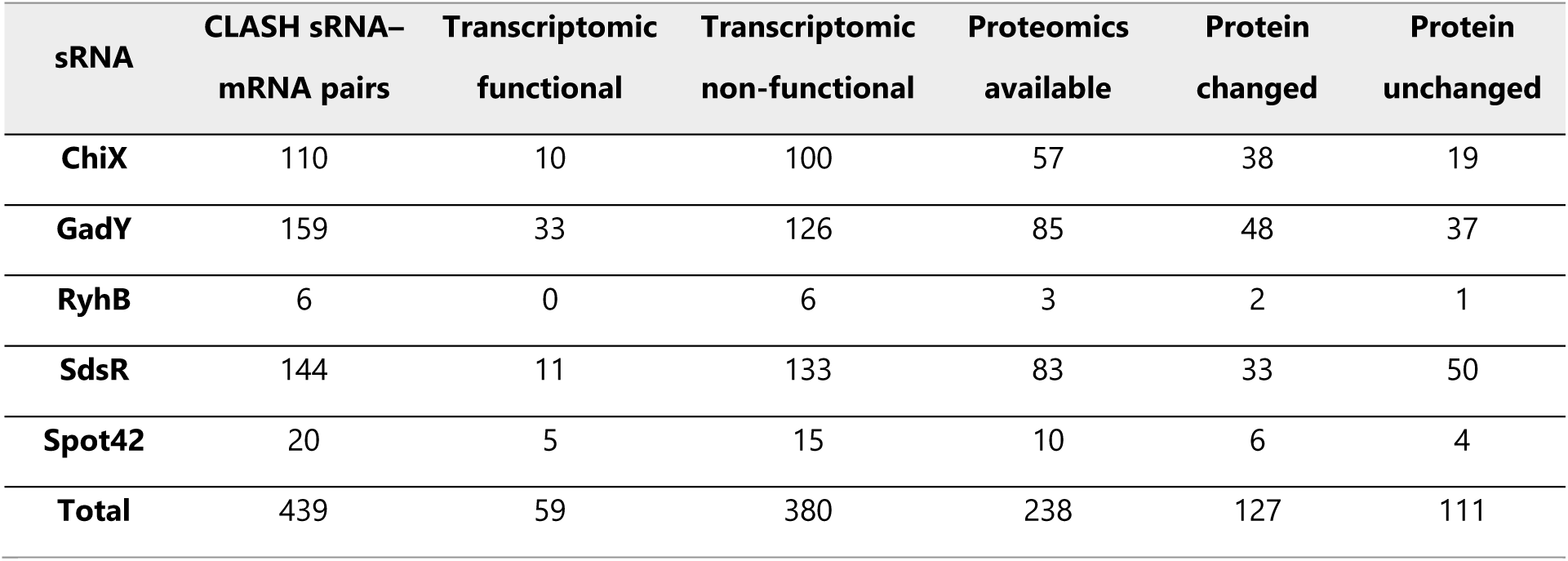
Distribution of CLASH-derived sRNA–mRNA interactions and functional outcomes across five hub sRNAs.

Functional annotation of this curated interaction set revealed that physical binding was not uniformly associated with regulatory output. At the transcriptomic level, only 59 of 439 interactions (13.4%) were associated with target mRNA changes, whereas 380 interactions (86.6%) showed no detectable RNA-level response despite being recovered by CLASH. Proteomic data were available for 238 pairs, of which 127 were associated with altered protein abundance and 111 showed no measurable protein-level response.

These results demonstrate that experimentally captured sRNA–mRNA interactions do not necessarily translate into functional regulation. Instead, regulatory consequences are layer-specific, with individual interactions producing transcript-level effects, protein-level effects, both, or neither under the conditions tested. This functionally annotated interaction set formed the basis for downstream modelling of transcriptomic response, proteomic response and hybrid-count abundance.

### Predictive Modelling of sRNA–mRNA Functional Outcomes via ML Pipelines

To determine whether the functional consequences of bacterial sRNA–mRNA interactions are intrinsically encoded within their molecular features, we implemented a comparative machine learning framework. We developed three independent pipelines to predict outcomes at the transcriptomic, proteomic, and hybrid-count levels using Hfq-CLASH-derived interaction data.

Systematic benchmarking of various feature selection and classification strategies revealed that logistic regression-based architectures consistently provided the highest discriminatory power. This is consistent with previous findings (36–39) in scenarios characterised by class imbalance and sample scarcity, where regularised logistic regression has been shown to be more robust and less prone to overfitting than higher-capacity or ensemble classifiers, which tend to require larger, better-balanced training sets to realise their performance advantage. For transcriptomic changes, the optimal configuration paired a Logistic Regression classifier with features identified via L2-regularized Logistic Regression. Prediction of proteomic shifts was best achieved using an LR-L2 classifier supported by RFE-LR feature selection. Finally, for hybrid-count classification, distinguishing high- from low-abundance chimeric ligation events, the top-performing model utilized both an LR-L2 classifier and LR-L2 feature selection.

All three optimised pipelines demonstrated discriminative performance above their respective no-discrimination baselines, with performance evaluated using ROC-AUC and PR-AUC. ROC-AUC was interpreted relative to the no-discrimination value of 0.5, whereas PR-AUC was interpreted relative to the positive-class prevalence within the corresponding held- out evaluation sets (Figure 5A, B).

**Figure 5.**
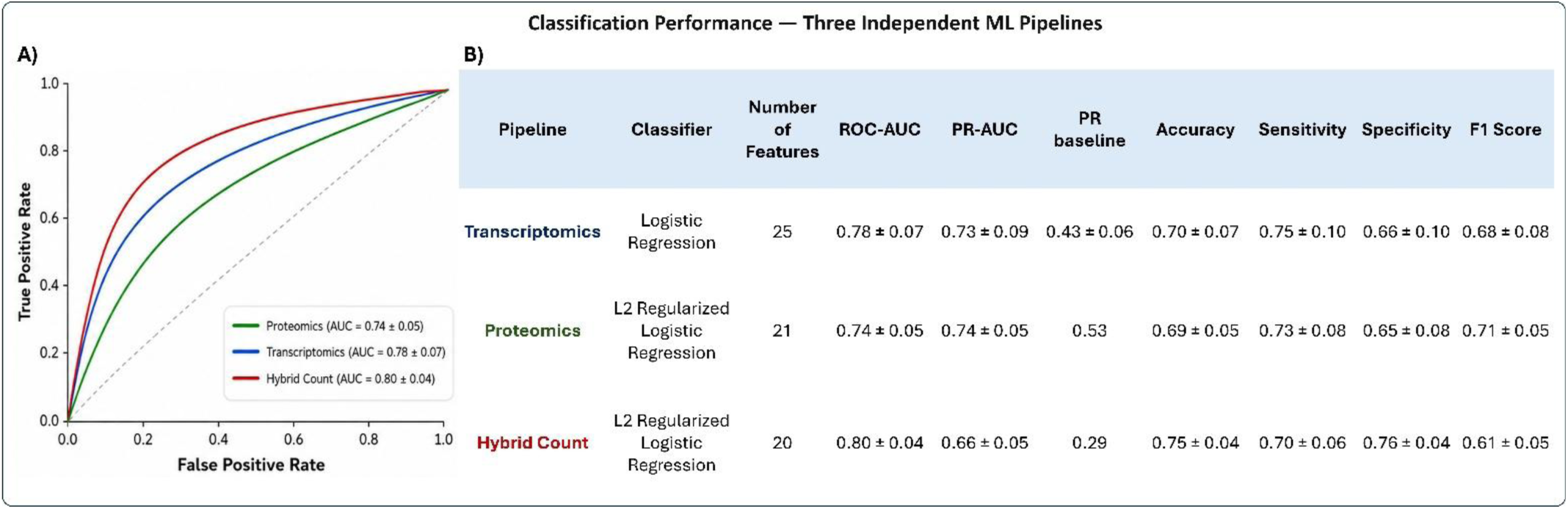
Comparative classification performance of the three independent ML pipelines. Receiver operating characteristic curves and summary performance metrics are shown for the optimised transcriptomics, proteomics and hybrid-count pipelines. ROC curves illustrate discrimination between functional and non-functional interactions for the transcriptomic and proteomic pipelines, and between high- and low-hybrid-count interactions for the hybrid-count pipeline. The accompanying table reports the best-performing classifier, number of retained features and mean ± SD values for ROC-AUC, PR-AUC (precision–recall area under the curve), PR baseline, accuracy, sensitivity, specificity and F1 score across held-out evaluation runs. ROC-AUC was interpreted relative to the no-discrimination baseline of 0.5, whereas PR-AUC was interpreted relative to the positive-class prevalence baseline for each pipeline.

The hybrid-count pipeline achieved the numerically highest ROC-AUC, with a ROC-AUC of 0.80 ± 0.04 and a PR-AUC of 0.66 ± 0.05, exceeding its positive-class prevalence baseline of 0.29. This model also showed the highest accuracy (0.75 ± 0.04) and specificity (0.76 ± 0.04). These results suggest that CLASH-derived interaction abundance, used here as a proxy for the physical frequency of sRNA–mRNA interactions, is tractable to classification from sequence, structural, thermodynamic and protein-occupancy features.

The transcriptomics pipeline, designed to predict target mRNA-level responses, achieved a ROC-AUC of 0.78 ± 0.07 and a PR-AUC of 0.73 ± 0.09. Because this pipeline used majority-class subsampling with out-of-bag evaluation, PR-AUC was interpreted relative to the positive-class prevalence within the held-out OOB sets (0.43 ± 0.06). The transcriptomics model also showed the highest sensitivity across the three biological readouts (0.75 ± 0.10), indicating that the selected molecular features were effective at recovering responsive interactions.

The proteomics pipeline achieved a ROC-AUC of 0.74 ± 0.05 and a PR-AUC of 0.74 ± 0.05 relative to a near-balanced PR baseline of 0.53. This model also achieved the highest F1 score (0.71 ± 0.05), showing a balanced precision–recall trade-off for detecting protein-level responses. Although the proteomics ROC-AUC was numerically lower than that of the transcriptomics and hybrid-count pipelines, this comparison needs to be placed in the context of the different biological endpoints, the number of labelled interactions available for each task and the measurement characteristics of the three datasets. Protein-level prediction may be influenced by both biological factors, including translational efficiency and protein turnover, and analytical factors, including proteome coverage, missingness, dynamic range and quantification variability. Together, these results indicate that experimentally observed sRNA–mRNA interactions contain molecular information predictive of hybrid-count abundance, transcript-level response and protein-level response. However, the predictive signal differs across readouts, supporting the view that physical interaction, mRNA response and protein response represent related but non-equivalent regulatory outcomes.

### Mapping Predictive Consistency across Individual sRNA–mRNA Interactions

To assess prediction stability beyond average performance metrics, we examined classification consistency for each sRNA–mRNA pair across the evaluation rounds in which it appeared in the held-out set (Fig. 6A–C). For each pipeline, we calculated the percentage of interactions correctly classified in at least 90% of rounds. Because the pipelines used different resampling designs, bootstrap out-of-bag evaluation for transcriptomics and repeated cross-validation for proteomics and hybrid count, these statistics were interpreted within each pipeline rather than compared directly across pipelines. Interaction-level prediction outputs for each pipeline are provided as Supplementary Files.

**Figure 6.**
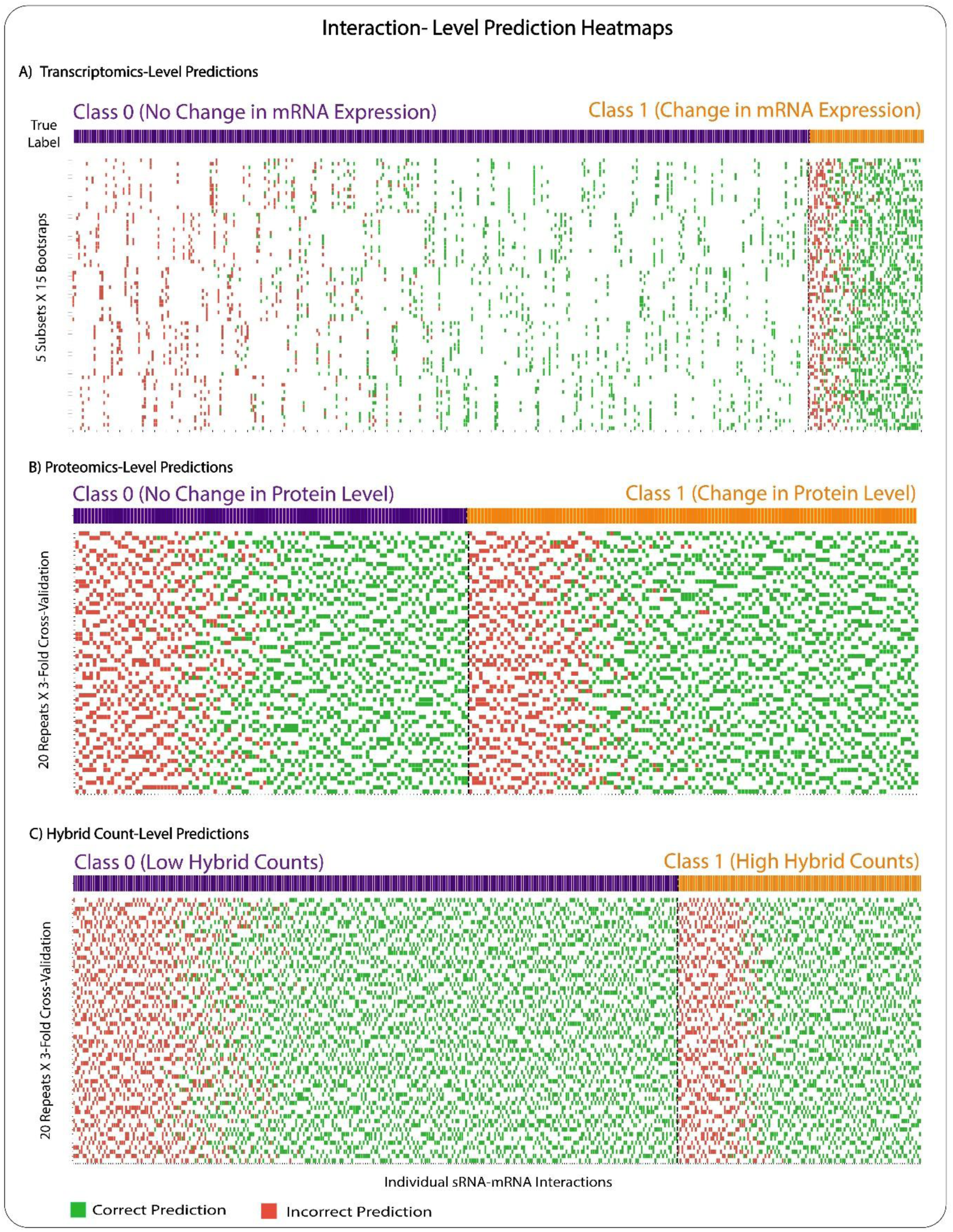
Interaction-level prediction consistency across biological readouts. Heatmaps show whether individual sRNA–mRNA interactions were correctly or incorrectly classified across repeated held-out evaluation rounds. Columns represent individual interactions, and rows represent bootstrap or cross-validation rounds. Green tiles indicate correct predictions, red tiles indicate incorrect predictions, and white tiles indicate rounds in which the interaction was not included in the held-out evaluation set. The top annotation bar indicates the true class, with class 0 representing non-responsive or low-abundance interactions and class 1 representing responsive or high-abundance interactions. (A) Transcriptomics-level predictions across 75 bootstrap out-of-bag rounds. (B) Proteomics-level predictions across 60 repeated cross-validation folds. (C) Hybrid-count predictions across 60 repeated cross-validation folds.

In the transcriptomics pipeline, 50.6% of interactions were correctly classified in at least 90% of rounds, with responsive interactions predicted at least as consistently as non-responsive interactions (55.9% versus 49.7%). In the proteomics pipeline, 57.1% of interactions met this consistency threshold, with higher stability for protein-responsive than non-responsive pairs (63.8% versus 49.5%). In the hybrid-count pipeline, 65.4% of interactions were consistently classified, with similar stability for low- and high-count classes (66.5% versus 62.7%). These interaction-level maps identify both reproducibly classified interactions and persistently ambiguous pairs, the latter representing candidates for targeted experimental follow-up.

### SHAP analysis identifies distinct feature-importance landscapes across pipelines

To investigate the molecular logic underlying our model predictions, we used SHAP analysis to identify the specific features driving classification at each biological layer (Figure 7A-C). This approach revealed that each pipeline relies on a unique set of molecular signatures, reflecting a transition from simple physical RNA pairing to more complex cellular regulation. In the transcriptomics pipeline, the model was primarily driven by the protein-binding context surrounding the interaction site (Figure 7A). High occupancy of the chaperone Hfq on both the sRNA and its target mRNA emerged as a strong predictor of a functional outcome, shifting predictions toward the responsive class. Conversely, high occupancy of the AR2 domain on the sRNA was a major driver for the non-responsive class, consistent with parallel work indicating that the AR2 sub-domain of RNase E recognises mRNA translation initiation regions and not sRNA (30). The influence of RNase E occupancy on the mRNA further highlights that transcriptomic changes are closely tied to the recruitment of the cellular machinery responsible for RNA decay.

**Figure 7.**
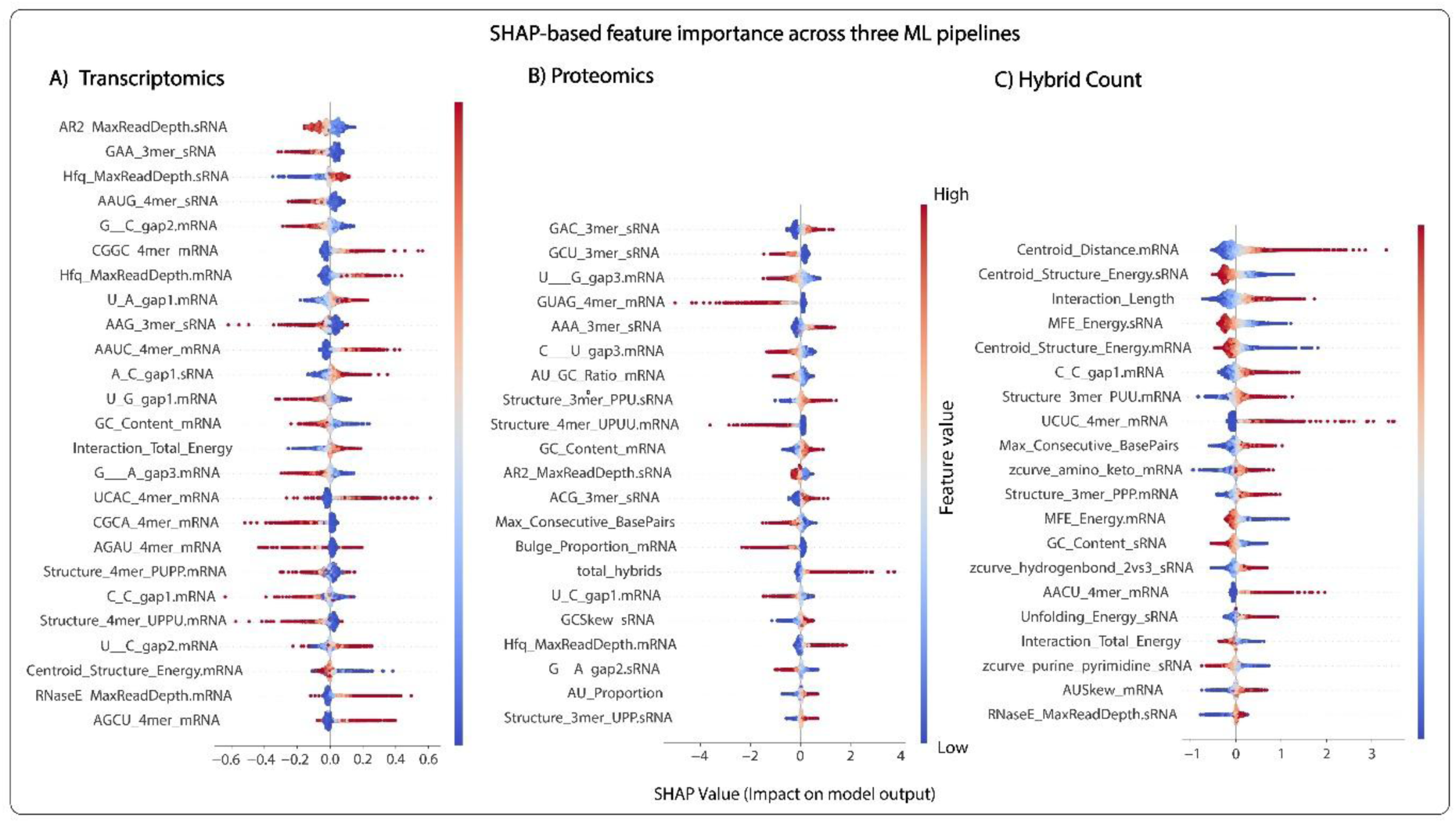
SHAP-based feature importance landscapes across regulatory layers. Summary plots illustrate the contribution of molecular features to the final model output for the (A) transcriptomics, (B) proteomics, and (C) hybrid-count pipelines. Each point represents an individual sRNA–mRNA interaction, with its position on the x-axis indicating the SHAP value (the magnitude and direction of the feature’s impact on the prediction). Positive SHAP values indicate a shift toward Class 1 (Responsive or High Abundance), while negative values indicate a shift toward Class 0 (Non-responsive or Low Abundance). Points are colored by the relative value of the feature (red = high, blue = low). Features are ranked vertically by their overall importance to the model. In panel A, the top features include protein-binding occupancy (e.g., Hfq and AR2). Panel B highlights the importance of "total hybrids" and specific sequence motifs. Panel C is dominated by biophysical and structural descriptors, such as centroid distance, interaction length, and thermodynamic energies. Vertical clusters of points indicate the distribution of feature impacts across the entire interaction dataset.

The proteomics model displayed a different landscape, relying heavily on interaction frequency and local sequence patterns (Figure 7B). A key driver in this pipeline was "total hybrids," which represents the overall read depth of interaction evidence in the Hfq-CLASH data. High interaction counts favoured the prediction of protein-level changes, indicating that the cumulative frequency of sRNA engagement is a robust indicator of eventual proteomic shifts. While Hfq occupancy on the mRNA remained an important feature, the model also placed significant weight on specific sequence motifs and gap features. This suggests that protein-level outcomes are influenced by a combination of the physical frequency of the interaction and the specific local sequence context of the target mRNA and sRNA.

The hybrid-count model was strongly influenced by biophysical and structural properties of the sRNA–mRNA pair (Figure 7C). The highest-ranking features included centroid distance and interaction length, with higher values generally shifting predictions toward the high-abundance class. Thermodynamic features, including MFE and centroid structure energy, also contributed to the model. Because these energy values are negative, lower values represent more stable predicted structures. In the SHAP analysis, lower MFE or centroid energy values generally shifted predictions toward the high-abundance class, indicating that thermodynamic stability can support recovery of abundant sRNA–mRNA hybrids. However, the model was not driven by stability alone; features related to centroid distance, interaction length, maximum consecutive base pairs, unfolding energy and local sequence composition also contributed to prediction. These results demonstrate that hybrid-count abundance reflects a combination of RNA stability, interaction geometry and target-site accessibility, rather than simply the strength of RNA–RNA binding. These shifting feature priorities across the three pipelines illustrate how the regulatory signal is transformed as it moves from the initial physical pairing event to measurable changes in the transcriptome and proteome.

### Shared coefficient signatures highlight common regulatory determinants

To identify molecular features with cross-layer predictive power, we compared standardized logistic regression (LR) coefficients for features selected by two or more pipelines (Figure 8). This comparative analysis revealed a core set of regulatory determinants that maintain consistent signals, alongside others that exhibit readout-dependent behaviour. Target-side Hfq occupancy emerged as a robust positive predictor for both transcriptomic (+0.060) and proteomic (+0.170) responses, reinforcing the established role of the Hfq chaperone in facilitating productive sRNA–mRNA engagement. In contrast, AR2 occupancy on the sRNA was systematically associated with the non-functional class across both transcriptomics (– 0.040) and proteomics (–0.140).

**Figure 8.**
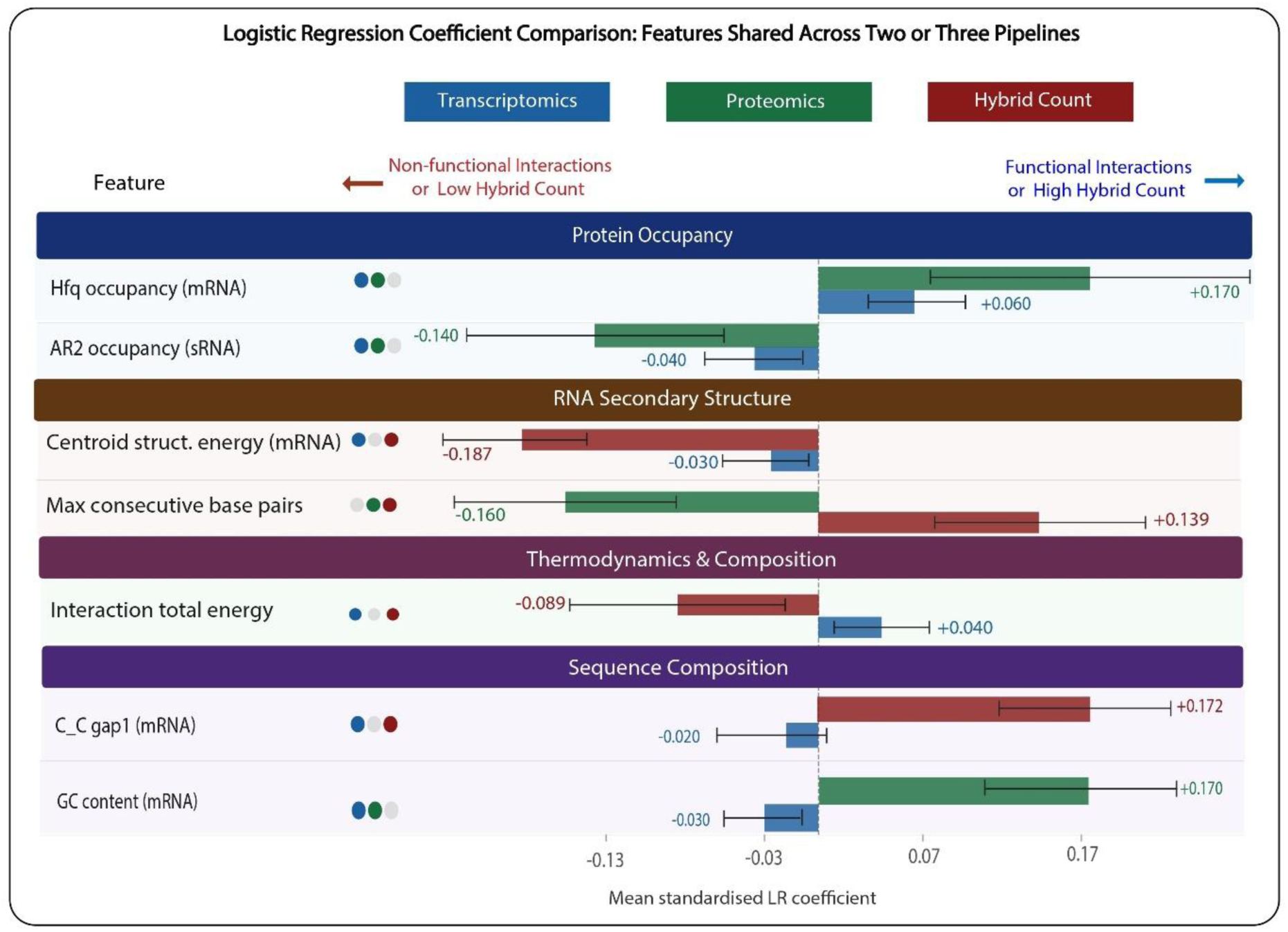
Comparative analysis of logistic regression coefficients for features shared across pipelines. Mean standardized logistic regression coefficients are shown for features selected by at least two pipelines: transcriptomics (blue), proteomics (green), and hybrid count (red). Positive values indicate an association with functional outcomes or high hybrid counts; negative values indicate associations with non-functional or low-abundance classes. Features are grouped by biological category, with colored dots indicating specific pipeline selection. Numerical labels highlight the mean coefficient values for primary regulatory determinants.

Beyond these consistent signals, several shared features displayed divergent directions, highlighting the distinct biophysical requirements for physical pairing versus functional outcome. For example, "max consecutive base pairs" was positively associated with high hybrid-count abundance (+0.139) but negatively correlated with proteomic shifts (–0.160). This divergence suggests that while extended, contiguous base-pairing supports the physical capture of chimeric fragments in CLASH experiments, it does not inherently guarantee protein-level regulation. Similarly, centroid structure energy of the mRNA showed negative coefficients in both the transcriptomics (–0.030) and hybrid-count (–0.187) models. Because these energy values are negative, this pattern suggests that lower, more stable mRNA structural energy values were associated with functional transcriptomic response and high hybrid-count abundance.

Readout-specific directions were also prominent among thermodynamic and sequence features. Interaction total energy showed opposite coefficient directions in the transcriptomics and hybrid-count models. In the transcriptomics model, its positive coefficient (+0.040) indicates that higher, less negative interaction energy values were associated with transcript-level response. By contrast, in the hybrid-count model, its negative coefficient (–0.089) indicates that lower, more negative interaction energy values were associated with high hybrid-count abundance. These findings support the idea that energetically favourable pairing may support physical recovery of abundant sRNA–mRNA hybrids, whereas transcriptomic response is not simply favoured by the strongest predicted duplex energy. Similarly, sequence-level features showed readout-specific effects; for example, GC content in the mRNA was negatively associated with transcriptomic response (– 0.030) but positively associated with proteomic response (+0.170), while C_C gap1 in the mRNA was negatively associated with transcriptomic response (–0.020) but positively associated with high hybrid-count abundance (+0.172). This pattern highlights that the predictive value of a molecular feature depends on the biological layer being modelled, where the same structural, thermodynamic or sequence property may support physical interaction recovery but have different implications for transcriptomic or proteomic outcomes.

### Readout-Specific Molecular Signatures Highlight the Complexity of sRNA-Mediated Regulation

To further resolve how distinct molecular feature categories contribute to each biological readout, we compared pipeline-unique coefficients across the three models (Figure 9). This analysis revealed that the transcriptomics model is heavily influenced by protein occupancy, specifically Hfq occupancy on the sRNA (+0.060) and RNase E occupancy on the mRNA (+0.050). These positive associations suggest that transcript-level changes are tightly coupled to the local assembly of the sRNA-chaperone-ribonuclease complex. Interestingly, the hybrid-count model also identified RNase E occupancy on the sRNA (+0.078) as a positive predictor, showing that ribonuclease-associated occupancy may enhance interaction capture or influence the abundance of detected chimeric fragments.

**Figure 9.**
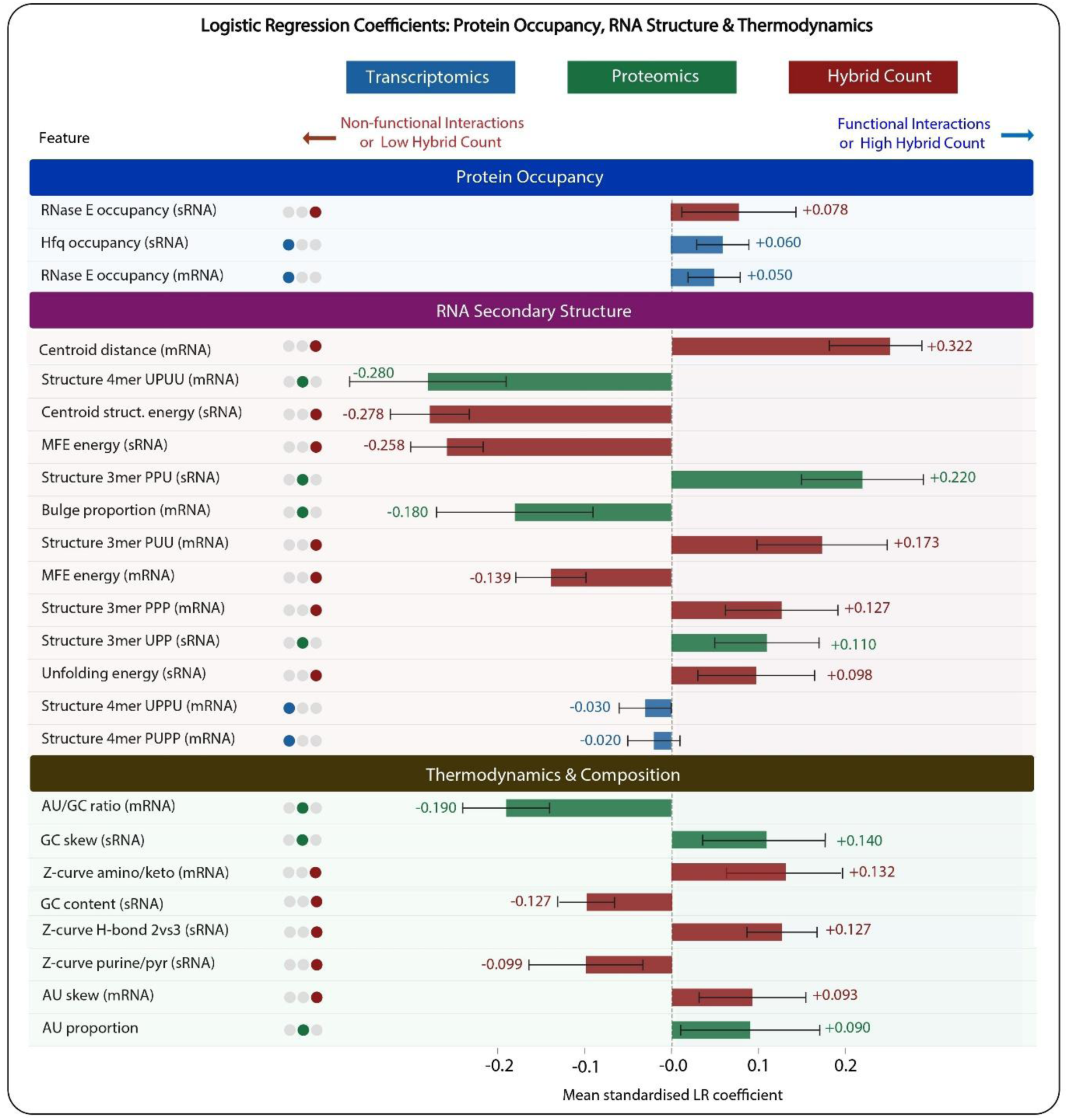
Comparison of pipeline-unique coefficients for protein occupancy, RNA structure, and thermodynamics. Standardized LR coefficients illustrate the predictive weight of features unique to each of the three modelling pipelines. Features are grouped into biologically interpretable categories: Protein Occupancy (derived from CRAC data), RNA Secondary Structure (including motif-based and energy descriptors), and Thermodynamics & Composition (e.g., Z-curve and AU-content metrics). The direction and magnitude of the bars highlight the distinct molecular logic driving transcriptomic (blue), proteomic (green), and hybrid-count (red) predictions. Colored dots denote the selecting pipeline for each feature.

RNA secondary structure and thermodynamic features contributed more to the prediction of proteomic and hybrid-count outcomes. In the proteomics model, local structural accessibility was a primary determinant, with negative coefficients observed for specific mRNA motifs like Structure 4mer UPUU (–0.280) and bulge proportion (–0.180). Conversely, specific sRNA structural motifs, such as Structure 3mer PPU (+0.220) and UPP (+0.110), were positively associated with protein-level shifts. These patterns may reflect differences in how specific local folding states and global RNA stability contribute to proteomic outcomes. In the hybrid-count pipeline, negative coefficients for sRNA centroid structure energy (–0.278) and MFE energy (–0.258) were observed, meaning that interactions with higher hybrid counts tended to involve sRNAs with lower, more stable predicted energies. This places sRNA structural stability among the features associated with hybrid abundance. This model instead favoured specific pairing motifs within the mRNA, including Structure 3mer PUU (+0.173) and PPP (+0.127), highlighting the potential relevance of both sRNA accessibility and the local structural context of the target to physical interaction recovery.

The contribution of sequence composition further differentiated the models, with the proteomics and hybrid-count pipelines showing a greater contribution from specific k-mer and gap features (Figure 10). The proteomics model included several sequence features with relatively larger coefficients, including negative weights for mRNA motifs such as GUAG 4mer (–0.360) and U_G gap3 (–0.270), alongside positive weights for sRNA motifs such as GAC 3mer and AAA 3mer (+0.240 each). In contrast, the transcriptomics model included a more compact set of sequence motifs with generally smaller coefficients, such as CGGC and AAUC 4mers in the mRNA. Compositional descriptors like the AU/GC ratio (–0.190) and AU proportion (+0.090) also informed proteomic predictions, whereas Z-curve descriptors provided unique signal for the hybrid-count model (Figure 9). These differences show that sequence and physicochemical properties contribute differently to prediction across transcriptomic, proteomic and interaction-level outcomes.

**Figure 10.**
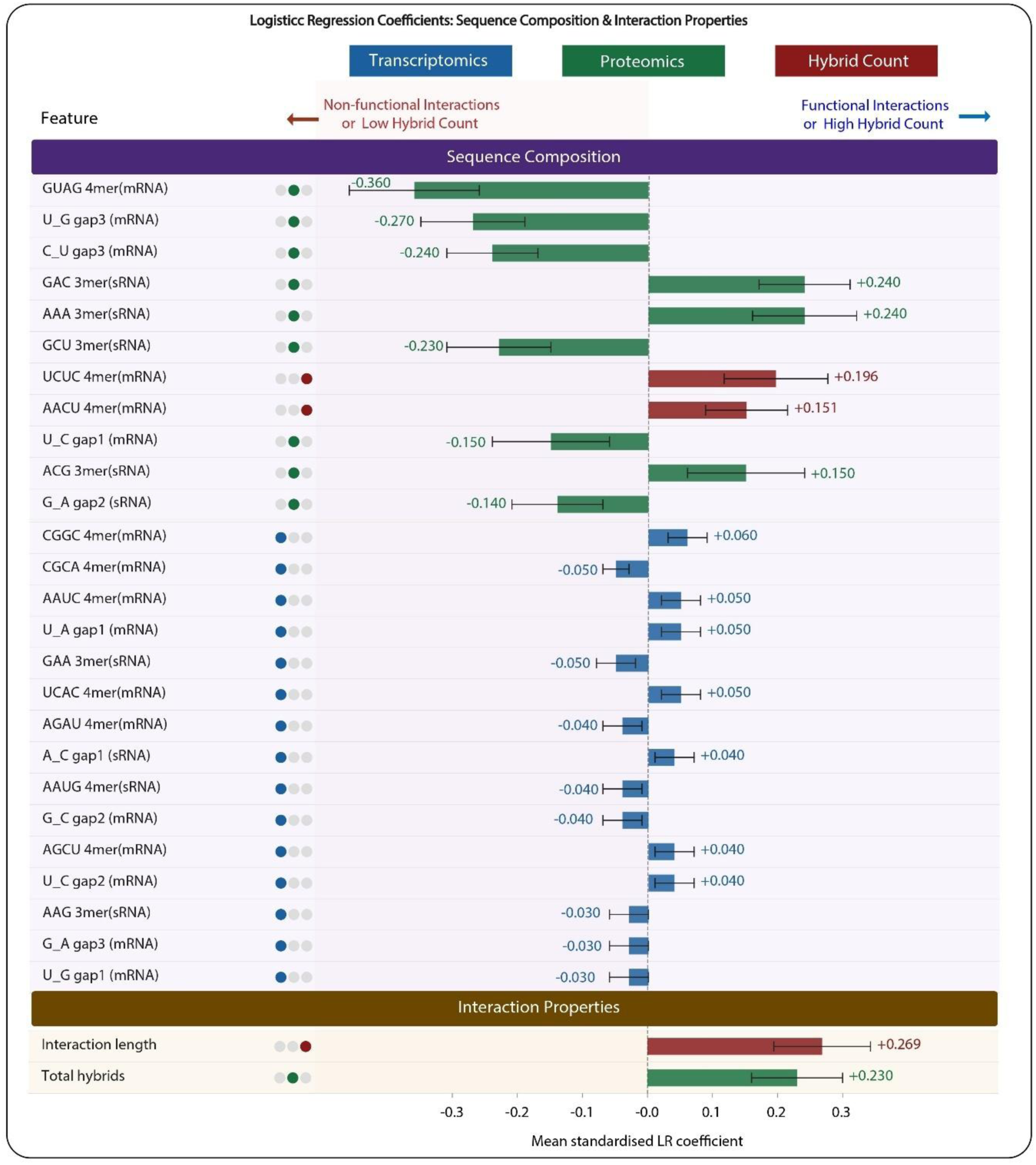
Comparison of pipeline-unique coefficients for sequence composition and interaction properties. Mean standardized logistic regression (LR) coefficients are shown for features selected specifically by the transcriptomics (blue), proteomics (green), or hybrid-count (red) pipelines. Features are categorized into Sequence Composition and Interaction Properties. Positive coefficients represent an association with functional regulatory outcomes or high hybrid-count abundance, while negative coefficients indicate an association with non-functional or low-abundance classes. Colored dots indicate the specific pipeline that utilized each feature.

The proteomics and hybrid-count models also differed in the interaction-related features associated with their predictions. In the proteomics model, Total Hybrids showed a positive coefficient (+0.230), linking greater hybrid recovery with the proteomic outcome. In the hybrid-count model, interaction length (+0.269) resulted in a positive coefficient, reflecting a contribution from structural properties of the interacting RNAs. These results show that while functional sRNA-mRNA regulation is detectable at multiple layers, the biological signal is fundamentally transformed as it moves from a physical pairing event (governed by geometry and structural accessibility) to a measurable functional change (governed by chaperone occupancy and sequence context).

## DISCUSSION

Transcriptome-wide RNA interaction mapping approaches, such as Hfq-CLASH and RIL-seq, have transformed the study of bacterial post-transcriptional regulation by revealing extensive networks of physical sRNA-mRNA interactions (18,40). However, these interaction maps also expose a central unresolved problem: hub sRNAs typically bind many more mRNA targets than the number that show detectable changes in transcript or protein abundance (14,18). Physical pairing is therefore an essential starting point for regulation, but it is not sufficient to define functional outcome. The present study addresses this gap by asking which molecular features distinguish experimentally detected interactions that produce transcriptomic or proteomic changes from those that remain functionally silent under the tested conditions.

By modelling transcriptomic response, proteomic response and hybrid-count abundance separately, we examined three related but non-equivalent layers of sRNA-mRNA regulation. The transcriptomics and proteomics models assessed downstream regulatory consequences, whereas the hybrid-count model captured variation in interaction recovery or abundance within the Hfq-CLASH dataset. The ability of all three readout-specific classifiers to discriminate their respective outcome classes above chance suggests that regulatory response and interaction abundance are associated with measurable properties of the interacting RNAs and their local protein-binding context. This supports a view in which functional outcome is shaped by the combined effects of protein occupancy, RNA sequence composition, secondary structure, thermodynamic features and interaction-level properties, rather than by physical contact alone.

The three pipelines showed distinct but informative performance profiles. The hybrid-count pipeline achieved the strongest overall discrimination, proposing that CLASH-derived interaction abundance is particularly tractable to prediction from RNA sequence, structure, thermodynamic and protein-occupancy features. This is consistent with hybrid count being the readout most proximal to the captured sRNA–mRNA interaction. The transcriptomics and proteomics pipelines also showed meaningful predictive performance, implying that molecular features associated with physical pairing retain information about downstream regulatory outcomes. However, the three pipelines model distinct biological endpoints and differ in labelled interaction availability and measurement characteristics, which should be considered when comparing their performance profiles. In particular, protein-level prediction may be influenced by both biological factors, such as translational efficiency, ribosome occupancy, protein stability and temporal regulation (1,17), and analytical factors, including proteome coverage, missingness, dynamic range and quantification variability.

Direct comparison with previous work is limited because, to our knowledge, this is the first study to model the functional outcome of experimentally captured bacterial sRNA–mRNA interactions across transcriptomic, proteomic and hybrid-count readouts. Most related computational studies instead focus on predicting physical sRNA–mRNA targets or interactions (41–43). Although these studies provide useful context for the achievable performance of sRNA target-prediction models, they address a different question from the present work. Rather than predicting whether two RNAs interact, our framework asks which molecular features distinguish experimentally observed interactions that are associated with downstream regulatory outcomes. In this context, the observed ROC-AUC and PR-AUC values support the conclusion that regulatory outcome is not random, but partially encoded in the biophysical and protein-binding context of each interaction.

A central finding across the models was the recurrent contribution of Hfq occupancy on the target mRNA, which was positively associated with functional outcome in both the transcriptomic and proteomic models. This is consistent with previous RIL-seq analyses showing that affected sRNA targets are recovered more frequently in sRNA-target duplexes and display higher Hfq occupancy than unaffected targets, whereas predicted duplex free energy alone shows only limited association with whether an interaction is recovered frequently or linked to downstream target regulation (16). These findings support a model in which productive regulation depends not only on base-pairing potential, but also on efficient target engagement with Hfq. Because Hfq is limiting, targets with higher Hfq occupancy may be more likely to compete successfully for sRNA pairing and downstream regulatory effects (16). Our results extend this model by showing that target-side Hfq occupancy remains predictive when analysed together with RNA sequence, structural, thermodynamic and protein-occupancy features.

RNase E-related occupancy provides mechanistic support for the idea that a subset of productive sRNA–mRNA interactions is coupled to RNA decay machinery. RNase E is the catalytic core of the bacterial RNA degradosome, a multienzyme complex involved in transcript turnover and RNA processing, and its C-terminal scaffold region helps organise interactions with degradosome components and regulatory factors (44). In Hfq-dependent regulation, sRNA pairing can promote RNase E-dependent cleavage and degradation of target mRNAs; this may occur after sRNA-mediated translational repression exposes the transcript to decay, or through more direct recruitment of RNase E to sRNA–mRNA complexes (45). Therefore, the positive contribution of RNase E occupancy in the models where it was retained as a predictor is biologically plausible and may indicate that functionally effective interactions are more likely when the target RNA is already connected to, or becomes recruited into, RNase E-associated decay pathways. However, because RNase E also participates in basal mRNA turnover, RNA maturation and broader RNA surveillance (46), this feature should be considered a marker of degradosome-linked RNA metabolism rather than a universal indicator of regulatory repression.

The AR2-associated signal should be interpreted more specifically than total RNase E occupancy. AR2 is an arginine-rich RNA-binding region within the intrinsically disordered C-terminal domain of RNase E; together with other RNA-binding regions, it contributes to RNA recognition and substrate engagement within the degradosome scaffold (12,47). The negative association between AR2 occupancy on the sRNA and functional transcriptomic or proteomic outcomes was unexpected, as AR2 has been proposed to help recruit Hfq-bound sRNA–mRNA complexes to the catalytic core of RNase E (47). However, recent work on the AR2 domain provides a possible explanation for this result. In that study, sRNAs were depleted relative to full-length RNase E, while AR2 binding was enriched at mRNA translation-initiation regions, consistent with AR2 engaging mRNAs before or during formation of productive sRNA–mRNA duplexes (30). The negative AR2 signal identified by the model therefore provides supporting evidence that AR2 occupancy carries predictive information distinct from total RNase E occupancy. Structural and thermodynamic descriptors in our models indicate that productive sRNA–mRNA regulation is not explained by sequence complementarity alone. Features such as centroid structure energy, MFE energy, bulge proportion, unfolding energy and maximum consecutive base pairs capture target-site accessibility, intrinsic RNA folding, the energetic cost of opening the interaction region and duplex continuity. This interpretation is consistent with established sRNA-target prediction frameworks, including TargetRNA2, IntaRNA and RNApredator, which incorporate accessibility, seed pairing, secondary structure and/or hybridisation energy to improve target identification beyond complementarity alone (41,43,48). Several structural features showed different coefficient directions across readouts, most plausibly because the same RNA property contributes differently to interaction recovery, mRNA-level regulation and protein-level output. Stronger or more continuous duplex formation, for instance, may favour recovery of RNA–RNA hybrids, while functional regulation at the transcriptomic or proteomic level depends on further layers of control, including correct positioning of the target site, occlusion or exposure of the ribosome-binding site, recruitment of RNase E and the resulting rate of RNA turnover. Hfq- and RNase E-CLASH studies support this distinction, showing that many detected sRNA–mRNA hybrids have strong base-pairing potential and enriched seed motifs, whereas only a subset produce measurable changes in mRNA abundance or translation (16,18).

The contribution of MFE and centroid structure energy features further suggests that RNA structural stability is relevant to sRNA–mRNA interaction outcomes, but its effect is context-dependent. Because these energy values are negative, lower values generally reflect more stable predicted structures. In our models, such features did not indicate a simple relationship in which either stability or accessibility alone determines function. Instead, stable structures may protect RNAs or maintain favourable conformations, while locally accessible regions remain important for initial sRNA docking and seed pairing. Hfq may further remodel or compact mRNA structure, bringing distant target sites into proximity with the sRNA and enabling kinetic selection of productive interaction sites (49). The structural signals detected in our models therefore likely reflect a balance between accessibility for initial pairing, duplex stability after pairing, and downstream regulatory consequences such as mRNA decay, stabilisation or translational control.

Sequence-composition features showed distinct, outcome-specific signatures across the transcriptomic, proteomic, and hybrid-count pipelines. We interpret these features as compact descriptors of local sequence context influencing pairing, accessibility, and protein-mediated recognition, rather than as deterministic regulatory motifs. In the proteomics model, positive coefficients for A-rich sRNA regions (e.g., AAA_3mer_sRNA) are biologically expected given that ARN-like sequences are recognized by the distal face of Hfq and can facilitate target annealing (50,51). Conversely, negative coefficients for certain mRNA-side sequence and gapped features likely reflect contexts less compatible with productive translational regulation, consistent with the principle that regulatory output depends heavily on target-site position, accessibility, and translational interference (52–54). This biophysical context is also reflected in the shifting influence of GC-content features, which highlight the trade-off between duplex stability and the energetic cost of opening structured target sites to permit pairing (55).

Similarly, the hybrid-count model was enriched for specific short sequence patterns alongside physical properties such as centroid distance and interaction length. Rather than representing downstream regulatory signals, these features likely reflect determinants of physical interactability and capture efficiency during Hfq-associated hybrid recovery. This interpretation agrees with Hfq-CLASH analyses showing that many recovered sRNA–mRNA hybrids have strong base-pairing potential and enriched complementary motifs, yet only a subset produces measurable mRNA-level changes (18). Overall, the sequence features in our models do not define a simple regulatory code; rather, they provide a local sequence framework through which structural accessibility, duplex geometry and Hfq/RNase E-associated processing shape the transition from physical pairing to transcriptomic and proteomic outcome.

The modelling strategy used here was designed to prioritise interpretability. Logistic regression, recursive feature elimination, SHAP-based attribution and coefficient analysis enabled each retained feature to be assigned a direction of association with the relevant outcome. This contrasts with higher-capacity approaches, such as graph-based or embedding-based models, which may improve predictive performance but are often less directly interpretable (21). For the present biological question, interpretability is essential because the goal is not only to classify interactions, but to generate mechanistic hypotheses about why some physical contacts become functional. The compact feature sets identified here provide candidate determinants that can be prioritised for targeted experimental validation.

Several limitations should be considered when interpreting these findings, and they also define important directions for future work. First, the analysis was restricted to 439 curated interactions involving five hub sRNAs, ChiX, GadY, RyhB, SdsR and Spot42, selected from a larger Hfq-CLASH interactome. Although this design enabled direct integration with matched transcriptomic and proteomic datasets, it remains unclear whether the same feature signatures generalise to other sRNAs, bacterial species, growth phases or environmental conditions. Second, the study focused on Hfq-associated interactions and therefore does not capture regulatory interactions mediated by other RNA-binding proteins or chaperones, such as ProQ or CsrA. Third, the transcriptomic and proteomic labels were defined using fold-change-based thresholds, which enabled supervised modelling but may not fully capture context-dependent or subtler regulatory effects. Proteomic predictions may also be influenced by analytical factors such as incomplete protein detection, missing values, dynamic range limitations and quantitative variability, as well as biological factors including translational efficiency and protein turnover. In addition, although the models identified informative associations between molecular features and regulatory outcomes, these relationships remain predictive rather than causal and require targeted experimental validation. Future studies should therefore extend this framework to larger and more diverse interactomes, including additional hub sRNAs, non-Hfq-associated interactions, other bacterial species and varied environmental conditions, to assess whether the readout-specific feature signatures observed here represent broader principles of bacterial sRNA-mediated regulation or are specific to the sRNAs and conditions examined. Integrating this interpretable feature-based framework with higher-capacity approaches, such as graph-based models or sequence-embedding methods, may also improve predictive performance.

The associations identified here generate testable hypotheses for targeted experimental follow-up. In particular, the negative relationship between sRNA-side AR2 occupancy and functional outcome could be examined using AR2 perturbation or domain-deletion approaches, combined with CRAC-based occupancy mapping for the five hub sRNAs studied here. Parallel AR2-focused work provides supporting evidence for this model, showing that sRNAs are depleted relative to full-length RNase E and that AR2 preferentially binds mRNA translation-initiation regions (30). Together, these analyses would help determine whether AR2-associated signals mark productive degradosome recruitment, non-productive engagement, or an intermediate state before functional sRNA–mRNA regulation.

## CONCLUSION

Overall, this study shows that the functional fate of an experimentally captured sRNA–mRNA interaction can be predicted, to a meaningful extent, from a combination of RNA sequence, structure, thermodynamic, interaction-level and protein-occupancy features. These findings support a model in which bacterial post-transcriptional regulation is not determined by physical RNA–RNA pairing alone, but emerges from the combined influence of the molecular properties of the interaction and the surrounding protein-binding context. In this framework, CLASH-detected interactions represent a spectrum of regulatory potential, with only a subset progressing from physical interaction to measurable transcriptomic or proteomic outcome.

## ACKNOWLEDGEMENTS

This research was supported by computational resources provided by the Katana high-performance computing cluster, managed by Research Technology Services at UNSW Sydney (https://research.unsw.edu.au/research-technology-services-restech). The authors acknowledge technical support from Monika Vesse. The graphical abstract was created in BioRender. Figure 1 was generated using GPT Image 2 and subsequently reviewed and modified by the authors. Generative AI tools were used to assist with language editing of the manuscript. All scientific content, interpretation and conclusions were generated by the authors, and the manuscript was thoroughly reviewed and verified by the authors.

## AUTHOR CONTRIBUTIONS

FS performed the investigation, feature extraction, model development and data analysis, generated the results and figures, and drafted the manuscript. FV and JT conceived and designed the study and supervised the research. FV supervised the development of the methodology. DGM and SA conducted the experiments. FV and JT provided funding and resources. All authors reviewed and revised the manuscript and approved the final version.

## DATA AVAILABILITY

Hfq-CLASH sequencing data is deposited at NCBI GEO under accession GSE343154. RNase E-CRAC and CRAC data for the AR2 sub-domain have been deposited at NCBI GEO under the accession GSE317719. RNA-seq data for sRNA overexpression studies have been deposited at NCBI GEO under accession GSE342672. The mass spectrometry proteomics data have been deposited to the ProteomeXchange Consortium via the PRIDE partner repository with the dataset identifier PXD082677.

## References

1. Tollerson R and Ibba M. Translational regulation of environmental adaptation in bacteria. J. Biol. Chem. 2020; 295: 10434. 10.1074/jbc.REV120.012742

2. Van Assche E, Van Puyvelde S, Vanderleyden J et al. RNA-binding proteins involved in post-transcriptional regulation in bacteria. Front. Microbiol. 2015; 6: 141. 10.3389/fmicb.2015.00141

3. Gottesman S. Post-transcriptional regulation and the bacterial response to stress. FASEB J. 2017; **31**: 22.1. 10.1096/fasebj.31.1_supplement.22.1

4. Chauvier A and Walter NG. Regulation of bacterial gene expression by non-coding RNA: it is all about time! *Cell Chem*. Biol. 2024; 31: 71. 10.1016/j.chembiol.2023.12.011

5. Ayenew Z, Eguale T, Bitew A et al. Current insights into the application of bacterial small RNAs in combating multidrug-resistant pathogens. Sci. Afr. 2026; 31: e03212. 10.1016/j.sciaf.2026.e03212

6. Sousa JP, Silva AFQ, Arraiano CM et al. Bacterial small RNAs: diversity of structure and function. RNA Technol. 2023; 14: 259–277. 10.1007/978-3-031-36390-0_12

7. He T, Ding Y, Sun Y et al. Advances in sRNA-mediated regulation of Salmonella infection in the host. Front. Cell. Infect. Microbiol. 2025; 15: 1503337. 10.3389/fcimb.2025.1503337

8. Storz G, Vogel J and Wassarman KM. Regulation by small RNAs in bacteria: expanding frontiers. Mol. Cell 2011; 43: 880. 10.1016/j.molcel.2011.08.022

9. Li X, Sun H, Yang X et al. sRNA centered signaling activates nitrate respiration and enhances Cronobacter sakazakii virulence in host environments. Nat. Commun. 2026; 17: 3373. 10.1038/s41467-026-70257-x

10. Cai LL, Xie YT, Hu HJ, et al. A small RNA, SaaS, promotes Salmonella pathogenicity by regulating invasion, intracellular growth, and virulence factors. Microbiol. Spectr. 2023; 11: e02938–22. 10.1128/spectrum.02938-22

11. Peer A and Margalit H. Accessibility and evolutionary conservation mark bacterial small-RNA target-binding regions. J. Bacteriol. 2011; 193: 1690. 10.1128/JB.01419-10

12. Alquethamy S, Lalaouna D and Tree JJ. What makes a small RNA work? Nucleic Acids Res.2025; 53: gkaf563. 10.1093/nar/gkaf563

13. Kavita K, de Mets F and Gottesman S. New aspects of RNA-based regulation by Hfq and its partner sRNAs. Curr. Opin. Microbiol. 2017; 42: 53. 10.1016/j.mib.2017.10.014

14. Melamed S, Peer A, Faigenbaum-Romm R et al. Global mapping of small RNA-target interactions in bacteria. Mol. Cell 2016; 63: 884–897. 10.1016/j.molcel.2016.07.026

15. Saliba AE, C Santos S and Vogel J. New RNA-seq approaches for the study of bacterial pathogens. Curr. Opin. Microbiol. 2017; 35: 78–87. 10.1016/j.mib.2017.01.001

16. Faigenbaum-Romm R, Reich A, Gatt YE et al. Hierarchy in Hfq chaperon occupancy of small RNA targets plays a major role in their regulation. Cell Rep. 2020; 30: 3127– 3138.e6. 10.1016/j.celrep.2020.02.016

17. Liu B, Chen H, Sheng K et al. Integrated transcriptomic and proteomic analyses reveal CsrA-mediated regulation of virulence and metabolism in Vibrio alginolyticus. Microorganisms 2025; 13: 1516. 10.3390/microorganisms13071516

18. Iosub IA, van Nues RW, McKellar SW et al. Hfq CLASH uncovers sRNA-target interaction networks linked to nutrient availability adaptation. eLife 2020; 9: e54655. 10.7554/eLife.54655

19. Moore KS and ’t Hoen PAC. Computational approaches for the analysis of RNA–protein interactions: a primer for biologists. J. Biol. Chem. 2019; 294: 1–9. 10.1074/jbc.REV118.004842

20. Xiao H, Yang X, Zhang Y et al. RNA-targeted small-molecule drug discoveries: a machine-learning perspective. RNA Biol. 2023; 20: 384. 10.1080/15476286.2023.2223498

21. Safari F, Tree JJ and Vafaee F. From nucleotides to numbers: a comprehensive review of RNA feature extraction methods for computational modelling. Brief. Bioinform. 2025; 26: bbaf701. 10.1093/bib/bbaf701

22. Wu W, Pang CNI, Tree JJ et al. Profiling the in vivo RNA interactome associated with the endoribonuclease RNase III in Staphylococcus aureus. Methods Enzymol. 2023; 692: 299– 324. 10.1016/bs.mie.2023.05.001

23. Tree JJ, Granneman S, McAteer SP et al. Identification of bacteriophage-encoded anti-sRNAs in pathogenic Escherichia coli. Mol. Cell 2014; 55: 199–213. 10.1016/j.molcel.2014.05.006

24. Tyanova S, Temu T and Cox J. The MaxQuant computational platform for mass spectrometry-based shotgun proteomics. Nat. Protoc. 2016; 11: 2301–2319. 10.1038/nprot.2016.136

25. Schoch CL, Ciufo S, Domrachev M et al. NCBI Taxonomy: a comprehensive update on curation, resources and tools. Database 2020; 2020: baaa062. 10.1093/database/baaa062

26. Perez-Riverol Y, Bandla C, Kundu DJ et al. The PRIDE database at 20 years: 2025 update. Nucleic Acids Res. 2025; 53: D543–D553. 10.1093/nar/gkae1011

27. Sy B, Wong J, Granneman S et al. High-resolution, high-throughput analysis of Hfq-binding sites using UV crosslinking and analysis of cDNA (CRAC). Methods Mol. Biol. 2018; 1737: 251–272. 10.1007/978-1-4939-7634-8_15

28. Mediati DG, Wong JL, Gao W et al. RNase III-CLASH of multi-drug resistant Staphylococcus aureus reveals a regulatory mRNA 3′UTR required for intermediate vancomycin resistance. Nat. Commun. 2022; 13: 3558. 10.1038/s41467-022-31177-8

29. Salgado H, Gama-Castro S, Lara P et al. RegulonDB v12.0: a comprehensive resource of transcriptional regulation in E. coli K-12. Nucleic Acids Res. 2024; 52: D255–D264. 10.1093/nar/gkad1072

30. Mediati DG, Alquethamy S, Jin C and Tree JJ. The intrinsically disordered AR2 domain of RNase E binds mRNA translation initiation regions. bioRxiv, 10.64898/2026.08.11.744331, 12 August 2026, preprint: not peer-reviewed.

31. Förstner KU, Vogel J and Sharma CM. READemption—a tool for the computational analysis of deep-sequencing-based transcriptome data. Bioinformatics 2014; 30: 3421– 3423. 10.1093/bioinformatics/btu533

32. Wu W, Pang CNI, Mediati DG et al. The functional small RNA interactome reveals targets for the vancomycin-responsive sRNA RsaOI in vancomycin-tolerant Staphylococcus aureus. mSystems 2024; 9: e00971–23. 10.1128/msystems.00971-23

33. Alvarez-Eraso KLF, Muñoz-Martínez LM, Alzate JF et al. Modulatory impact of the sRNA Mcr11 in two clinical isolates of Mycobacterium tuberculosis. Curr. Microbiol. 2022; 79: 39. 10.1007/s00284-021-02733-0

34. Hoang TM, Huang W, Gans J et al. The heme-responsive PrrH sRNA regulates Pseudomonas aeruginosa pyochelin gene expression. mSphere 2023; 8: e00392–23. 10.1128/msphere.00392-23

35. Shrestha S, Awasthi D, Chen Y et al. Simultaneous carbon catabolite repression governs sugar and aromatic co-utilization in Pseudomonas putida M2. Appl. Environ. Microbiol. 2023; 89: e00852–23. 10.1128/aem.00852-23

36. Shahri NHNBM, Lai SBS, Mohamad MB et al. Comparing the performance of AdaBoost, XGBoost, and logistic regression for imbalanced data. Math. Stat. 2021; 9: 379–385. 10.13189/ms.2021.090320

37. Vijayan A, Fatima S, Sowmya A et al. Blood-based transcriptomic signature panel identification for cancer diagnosis: benchmarking of feature extraction methods. Brief. Bioinform. 2022; 23: bbac315. 10.1093/bib/bbac315

38. Li D, Heffernan K, Koch FC et al. Discovery of plasma lipids as potential biomarkers distinguishing breast cancer patients from healthy controls. Int. J. Mol. Sci. 2024; 25: 11559. 10.3390/ijms252111559

39. Rahman HAA, Wah YB, He H, et al. Comparisons of ADABOOST, KNN, SVM and logistic regression in classification of imbalanced dataset. Commun. Comput. Inf. Sci. 2015; 545: 54–64. 10.1007/978-981-287-936-3_6

40. Mizrahi SP, Elbaz N, Argaman L et al. The impact of Hfq-mediated sRNA-mRNA interactome on the virulence of enteropathogenic Escherichia coli. Sci. Adv. 2021; 7: eabi8228. 10.1126/sciadv.abi8228

41. Kery MB, Feldman M, Livny J et al. TargetRNA2: identifying targets of small regulatory RNAs in bacteria. Nucleic Acids Res. 2014; 42: W124. 10.1093/nar/gku317

42. Tjaden B. TargetRNA3: predicting prokaryotic RNA regulatory targets with machine learning. Genome Biol. 2023; 24: 276. 10.1186/s13059-023-03117-2

43. Busch A, Richter AS and Backofen R. IntaRNA: efficient prediction of bacterial sRNA targets incorporating target site accessibility and seed regions. Bioinformatics 2008; 24: 2849. 10.1093/bioinformatics/btn544

44. Worrall JAR, Górna M, Crump NT et al. Reconstitution and analysis of the multienzyme Escherichia coli RNA degradosome. J. Mol. Biol. 2008; 382: 870. 10.1016/j.jmb.2008.07.059

45. Prévost K, Desnoyers G, Jacques JF et al. Small RNA-induced mRNA degradation achieved through both translation block and activated cleavage. Genes Dev. 2011; 25: 385. 10.1101/gad.2001711

46. Bandyra KJ and Luisi BF. RNase E and the high-fidelity orchestration of RNA metabolism. Microbiol. Spectr. 2018; 6: RWR-0008-2017. 10.1128/microbiolspec.RWR-0008-2017

47. Sinha D and De Lay NR. Target recognition by RNase E RNA-binding domain AR2 drives sRNA decay in the absence of PNPase. Proc. Natl. Acad. Sci. U.S.A. 2022; 119: e2208022119. 10.1073/pnas.2208022119

48. Eggenhofer F, Tafer H, Stadler PF et al. RNApredator: fast accessibility-based prediction of sRNA targets. Nucleic Acids Res. 2011; 39: W149. 10.1093/nar/gkr467

49. Hoekzema M, Romilly C, Holmqvist E et al. Hfq-dependent mRNA unfolding promotes sRNA-based inhibition of translation. EMBO J. 2019; 38: e101199. 10.15252/embj.2018101199

50. Link TM, Valentin-Hansen P and Brennan RG. Structure of Escherichia coli Hfq bound to polyriboadenylate RNA. Proc. Natl. Acad. Sci. U.S.A. 2009; 106: 19292–19297. 10.1073/pnas.0908744106

51. Updegrove TB, Zhang A and Storz G. Hfq: the flexible RNA matchmaker. Curr. Opin. Microbiol. 2016; 30: 133. 10.1016/j.mib.2016.02.003

52. Azam MS and Vanderpool CK. Translational regulation by bacterial small RNAs via an unusual Hfq-dependent mechanism. Nucleic Acids Res. 2018; 46: 2585–2599. 10.1093/nar/gkx1286

53. Djapgne L and Oglesby AG. Impacts of small RNAs and their chaperones on bacterial pathogenicity. Front. Cell. Infect. Microbiol. 2021; 11: 604511. 10.3389/fcimb.2021.604511

54. Bandyra KJ, Said N, Pfeiffer V et al. The seed region of a small RNA drives the controlled destruction of the target mRNA by the endoribonuclease RNase E. Mol. Cell 2012; 47: 943–953. 10.1016/j.molcel.2012.07.015

55. Raden M, Müller T, Mautner S et al. The impact of various seed, accessibility and interaction constraints on sRNA target prediction: a systematic assessment. BMC Bioinformatics 2020; 21: 15. 10.1186/s12859-019-3143-4

