## Supplementary material for "Molecular Determinants of Functional Bacterial sRNA–mRNA Interactions Revealed by Integrating RNA Interactomes and Interpretable Machine Learning": Supplementary_ MATERIAL AND METHODS.docx

**Experimental Procedures**

***E. coli* MG1655 Hfq-CLASH**

CLASH was performed as previously described (1) with some modifications specific for *E. coli*. MG1655 WT and Hfq-HTF (dual-affinity-tagged HTF strain) (2) were used to inoculate 400 mL of pre-warmed liquid LB and grown at 37°C to an OD_600nm_ 2.0 with 160 rpm shaking. Cultures were crosslinked with 1800 mJ of UV-C (Vari-X-Link, UV03) and immediately harvested by vacuum onto a membrane filter. Each culture condition was performed in triplicate. One mL of lysis buffer (50 mM Tris-HCl (pH 7.8), 1.5 mM MgCl_2_, 150 mM NaCl, 0.1% NP-40, 5mM b-mercaptoethanol and 1 tablet of “cOmplete” EDTA-free protease inhibitor (Roche) per 50 mL of buffer) and 2 V of 0.1 mm zirconia beads were added to each cell pellet and vortexed for 2 x 40 sec intervals using the FastPrep-24 5G (MP Biomedicals). Dry ice was used to ensure cell pellets stayed chilled during intervals. Cell debris was centrifuged (4,500 *g* for 20 min) and the clarified lysate was transferred to 1.5 mL microcentrifuge tubes and further clarified at 16,000 *g* for 20 min. Supernatants were added to equilibrated M2 anti-FLAG resin (Sigma) and incubated at 4°C with gentle rotation for 16 h. The resin was washed twice in 5 mL TNM1000 buffer (50 mM Tris-HCl (pH 7.8), 1 M NaCl, 0.1% NP-40, 5mM b-mercaptoethanol) and twice in 10 mL TNM150 (50 mM Tris-HCl (pH 7.8), 150 mM NaCl, 0.1% NP-40, 5mM b-mercaptoethanol). The protein-bound resin was resuspended in 500 mL of TNM150 and incubated with 50 U of GST.TEV protease at 24°C with rotation for 2 h. Eluates were collected by filtration through a Bio-spin chromatography column (Bio-Rad) and incubated with 0.15 U of RNace-IT (Agilent) at 20°C for 5 min. The digestion stopped by the addition of 0.4 g guanidine-HCl, 300 mM NaCl and 10 mM imidazole. Eluates were added to 150 mL magnetic Ni-NTA resin (Thermo) equilibrated with wash buffer 1 (4 M guanidine-HCl, 50 mM Tris-HCl (pH 7.8), 300 mM NaCl, 0.1% NP-40, 5mM b-mercaptoethanol) and incubated at 4°C with gentle rotation for 16 h. Ni-NTA resin was washed twice with 1 mL wash buffer 1 and 3x with 1 mL PNK buffer (50 mM Tris-HCl (pH 7.8), 10 mM MgCl_2_, 0.1% NP-40, 5mM b-mercaptoethanol). The resin was resuspended in 80 mL NP-PNK buffer (50 mM Tris-HCl (pH 7.8), 10 mM MgCl, 5mM b-mercaptoethanol) containing 8 U of alkaline phosphatase (Promega) and 80 U of recombinant RNasIN (Promega) and incubated at 37°C with gentle rotation for 1 h. The magnetic Ni-NTA resin was washed once with 1 mL wash buffer 1 and 3x with 1 mL PNK buffer, and resuspended in 80 mL NP-PNK buffer containing 20 U of T4 polynucleotide kinase (NEB), 20 U of RNasIN (Promega) and 30 mCi ^32^P-ATP and incubated at 37°C with rotation for 50 min. Cold 1 mM ATP was spiked in and the reaction allowed to proceed for another 10 min to ensure all 5’ end phosphates are attached. Ni-NTA resin was again washed once with 1 mL wash buffer 1 and 3x with 1 mL PNK buffer and then resuspended in 80 mL of NP-PNK buffer containing 40 U of T4 RNA ligase I (NEB), 1 mM ATP, 20 U of RNasIN (Promega) and 100 pM of an L5 index barcoded 5’ linker (IDT). The ligation reaction was incubated at 16°C with gentle rotation for 16 h. The magnetic Ni-NTA resin was washed once with 1 mL wash buffer 1 and 3x with 1 mL PNK buffer, and resuspended in 60 mL NP-PNK buffer containing 40 U of T4 RNA ligase I (NEB), 20 U of RNasIn (Promega) and 80 pM of 3’ linker App-PE (IDT). The ligation reaction was incubated at 16°C with gentle rotation for 16 h. Ni-NTA resin was washed once with 1 mL wash buffer 1 and 3x with 1 mL wash buffer 2 (50 mM Tris-HCl (pH 7.8), 50 mM NaCl, 0.1% NP-40, 5mM b-mercaptoethanol, 10 mM imidazole). ^32^P-labelled Hfq-RNA complexes were eluted in 200 mL elution buffer (50 mM Tris-HCl (pH 7.8), 50 mM NaCl, 0.1% NP-40, 5mM b-mercaptoethanol, 250 mM imidazole) at 8°C with rotation for 2x 15 min repeats and precipitated with TCA for 1 h. The pellets were washed 2x with acetone and resolved on a NuPAGE 4-12% gradient Bis-Tris PAGE gel (Invitrogen) run at 4°C in 1x NuPAGE running buffer for 1 h at 150 V. ^32^P-labelled Hfq-RNA complexes were visualised by autoradiography, gel excised and recovered by incubating the fragmented gel pieces in 500 mL of wash buffer 3 (50 mM Tris-HCl (pH 7.8), 50 mM NaCl, 0.1% NP-40, 5mM b-mercaptoethanol, 1% SDS, 5 mM EDTA) and 100 mg of proteinase K at 55°C for 2 h. Remaining gel pieces were removed by filtration through a Bio-spin chromatography column (Bio-Rad) and RNA eluates were phenol:chloroform purified and ethanol precipitated overnight at -80°C. Reverse transcription was performed using the RT_PE_reverse oligo and SuperScript IV (Invitrogen) according to the manufacturer’s instructions. cDNA was incubated with RNase H (Thermo) at 37°C for 15 min and amplified using Phusion Hot Start Flex DNA Polymerase (NEB) according to the manufacturer’s instructions. A combination of either 10 pM BC1, BC2 or BC3 oligonucleotides were used with 10 pM of the P5 oligonucleotide to alternate the unique barcoded indexes. The cDNAs were amplified for 22 cycles. cDNA libraries were separated on a 1.5% metaphor agarose gel and amplicons excised and purified using the MiniElute gel extraction kit (Qiagen). Libraries were sequenced on a NovaSeq 6000 platform generating 150 bp paired-ended reads (PE150) (Novagene, Singapore).

**References**

1.	Wu W, Pang CNI, Tree JJ, Mediati DG. Profiling the in vivo RNA interactome associated with the endoribonuclease RNase III in Staphylococcus aureus. Methods Enzymol. 2023 Jan 1;692:299–324. doi:10.1016/bs.mie.2023.05.001 PubMed PMID: 37925184.

2.	Tree JJ, Granneman S, McAteer SP, Tollervey D, Gally DL. Identification of Bacteriophage-Encoded Anti-sRNAs in Pathogenic Escherichia coli. Mol Cell. 2014 Jul 17;55(2):199–213. doi:10.1016/j.molcel.2014.05.006 PubMed PMID: 24910100.
